# Ultrasonic Modulation of Astrocytic and Neuronal Calcium Dynamics in Mouse Cortex

**DOI:** 10.1101/2025.03.03.641164

**Authors:** Yu Yong, Hao Jiang, Chaofeng Qiao, Zhuoyan Liu, Xiaohan Zhou, Yufeng Zhou, Fenfang Li

## Abstract

**Background:** Astrocytes are abundant in the brain and their calcium signaling is reported to have an important effect on neuronal activity in both physiological and pathological conditions. Low-frequency focused ultrasound (FUS) has recently emerged as a powerful noninvasive neuromodulation approach, yet its impact on astrocyte calcium dynamics in different brain states *in vivo* is poorly understood.

**Objective:** This study aimed to elucidate the effects of non-thermal FUS on astrocyte calcium dynamic with *in vivo* cellular-resolution and cell-type-specific recording and identify whether the influences of FUS on cortical astrocytes and neurons are distinctive and state dependent.

**Methods:** Here we combined a customized 0.521MHz FUS transducer with two-photon microscopy, allowing simultaneous single-cell resoultion imaging and FUS stimulation at intensities of 0.91 or 1.5 W/cm^2^ to examine astrocyte and neuronal calcium responses in somatosensory cortex of both awake and lightly anesthetized mice. Functional clustering analysis was performed to identify calcium response activated or inhibited subpopulations.

**Results:** In awake mice, FUS significantly enhanced the amplitude, frequency, and temporal integral of astrocyte calcium transients, while suppressing neuronal calcium activity and reducing the proportion of activated neuronal subpopulations. In contrast, lightly anesthetized mice displayed a blunted yet increased astrocyte response and negligible neuronal modulation under FUS, suggesting that baseline suppression from anesthesia partially masks FUS effects.

**Conclusions:** Our study demonstrated that FUS elicited distinctive, state-dependent effects on cortical astrocytes and neurons, highlighting astrocytes as previously underappreciated targets of ultrasound neuromodulation. These findings will pave the way for FUS-based therapies targeting astrocyte–neuron interactions in conditions involving abnormal brain excitability.

**Highlights:**

- Low duty cycle low frequency focused ultrasound induced negligible heat effects.
- Ultrasound markedly enhanced astrocyte calcium transients in awake mice.
- Ultrasound suppressed neuronal calcium activity in awake mice.
- The effects of ultrasound on both cell types were reduced under light anesthesia.

## Introduction

Neuromodulation techniques are indispensable in advancing our understanding of the complex brain processes and the pathophysiology of neurological disorders^1^. Although electrical brain stimulation modalities, such as deep brain stimulation, spinal cord stimulation, or motor cortex stimulation can be effective, they often require the surgical implantation of electrodes in the brain, accompanied by inevitable risks^2,3^. Non-invasive strategies like transcranial direct current stimulation and repetitive transcranial magnetic stimulation offer safer alternatives but have limited depth of penetration and lacking spatial resolution^4–6^. In contrast, focused ultrasound (FUS) stimulation has emerged as a promising non-invasive method capable of modulating neuronal activity with higher spatial temporal precision^7^. At low intensities, FUS can penetrate the skull and dura to influence neural connectivity and neurotransmission, making it attractive for therapeutic applications^8–11^.

Although accumulating evidence suggests that FUS can either evoke or suppress neural activity depending on the brain subregions and stimulation parameters^12,13^, much less is known about its effects on astrocyte calcium dynamics. Astrocytes are not merely supportive cells; they are actively involved in synaptic function and brain network homeostasis with their extensive connections and calcium-based excitablity^14–16^. Some *in vitro* studies have shown that low-intensity, low-frequency ultrasound opens the transient receptor potential channel A1 (TRPA1) in cultured astrocytes, leading to calcium influx and glutamate or GABA-dependent synaptic transmission^17–19^. Moreover, the elevated intracellular calcium, enhanced expression of neurotrophic factors, and the release of exosomes have been observed in cultured astrocytes after ultrasound stimulation^20–22^. However, the precise nature and dynamics of FUS-modulated astrocyte calcium at cellular resolution *in vivo* remain largely unexplored. Unraveling this gap is essential for understanding the full scope of FUS-mediated brain stimulation.

Astrocytes can regulate neuronal activity through calcium-dependent signaling^15,23^. For instance, it was found that calcium influx in astrocytes triggered rapid endocytosis of the GABA transporter and increased synaptic GABA levels, contributing to the neuronal inhibition in fruit flies^24^. Rodent hippocampal studies have revealed that calcium activation in astrocytes induce adenosine triphosphate (ATP) and elevated tonic GABA release, mediating synaptic depression and suppression of hippocampal circuit function^25–28^. Additionally, in Alzheimer’s disease mouse model, chemogenetic recovery of the decreased astrocyte calcium mitigated neuronal hyperactivity, supporting that astrocyte calcium signaling negatively regulates neuronal activity^29^. These findings underscore the importance of astrocyte calcium dynamics in shaping neuronal network activity, yet how FUS modulates astrocyte calcium dynamics, and the relevant neuronal activity change *in vivo* remains unclear.

The effects of FUS on neuronal calcium dynamics and activity are well documented. In small animals like mice and rats, low-intensity FUS has been shown to increase spike firing in cerebellum, hippocampus, and primary somatosensory cortex^9,30–32^. Studies also showed that FUS could increase the activity of interneurons while suppressing excitatory neurons in hippocampus or suppress neural activity in visual cortex, motor cortex and thalamus at different foundamental frequencies and pulse repetition frequencies (PRF)^33–36^. Adjusting the stimulation modules of FUS allows bidirectional control of neural activity in both animals and human^37–41^. However, whether FUS elicits similar or different responses in astrocytes and neurons is unknown.

To address this gap, we investigated the effects of FUS on the astrocytic and neuronal calcium dynamics in both awake and anesthetized mice. We also explored the application of focused ultrasound with 0.521MHz fundamental frequency at 1.5 and 0.91 W/cm^2^ intensity. By integrating a ring-shaped focused ultrasound transducer with *in vivo* two-photon calcium imaging, we recorded and analyzed the activity of astrocytes and neurons at cellular resolution in various conditions. Our detailed characterization of the astrocytic and neuronal activity in response to FUS with different stimulation intensities and brain states provides important insights for further development of ultrasound-based brain stimulation techniques and therapies.

## Methods

### 1. Stereotaxic injection and cranial window surgery

The wild-type C57BL/6 mice were purchased from GemPharmatech. Mice (8–12 weeks old) were kept at 37 °C using a heating pad during all surgical and imaging procedures. Surgery was performed under isoflurane anesthesia (1.5–2%). Following shaving of the head, disinfection of the skin, an incision was made to expose the skull. The skull was dried by removing all bone-attached membrane and fat tissue. Mice were then placed into a stereotactic injection apparatus.

A hole was drilled in the skull to inject rAAV-GfaABC1D-GCaMp6f-WPRE-hGH polyA virus (PT-2560, BrainVTA) and/or rAAV-hSyn-NES-jRGECO1a-WPRE-hGH polyA virus (PT-1593, BrainVTA) for the respective detection of astrocytic and neuronal calcium dynamics in barrel cortex (−1.7 mm anterior-posterior, +2.65 mm medial-lateral and −0.85 mm dorsal-ventral, relative to bregma) using a beveled 33-gauge Hamilton needle at a rate of 50 nl/min. A craniotomy over the right whisker barrel cortex, including the stereotactic injection site, was prepared immediately afterwards. Two small self-tapping bone screws (M1) were inserted into the skull for anchoring of the dental cement and head bar. A custom-designed head bar that included an imaging chamber was positioned over the craniotomy and firmly affixed to the skull with superglue and dental cement. The exposed cortex was covered by a round glass coverslip (4 mm in diameter, 0.15 mm thick), which was affixed to the skull and sealed with Flow-It ALC composite. Mice were administered meloxicam and allowed to recover. Imaging experiments were performed 21–30 days later. All procedures involving mice were approved by the Animal Care and Use Committees of Shenzhen Bay Laboratory.

### 2. Focused ultrasound stimulation and characterization

Ultrasound waves were generated using a 0.521 MHz ring-shaped transducer (the emitting surface with a concentric spherical curvature, see Fig. S1) for mouse brain stimulation. All stimulations were performed using two connected function generators (Rigol DG1022 and DG972) through a power amplifier (E&I A075). The waveforms were monitored through an oscilloscope (Teledyne LeCroy 4104HD). The transducer with customized coupling cone filled with de-gassed ultrasound gel was installed on the Olympus 25× objective, aligning the focal plane of two-photon laser with ultrasound (see Fig. S1). Mice were stimulated with or without focused ultrasound for 1min under the 5% duty cycle, 0.91 W/cm^2^ spatial-peak pulse-average intensity (I_SPPA_) or 10% duty cycle, 1.5 W/cm^2^ I_SPPA_. The acoustic field and output power of the ultrasound transducer were measured with a submersible hydrophone (HNR0500, ONDA) placed beneath the coverslip used for cranial window in a water tank. For more details of the acoustic field measurement, please refer to the Supplemental Materials.

### 3. *In vivo* two-photon imaging

An Olympus multiphoton microscope (FVMPE-RS) equipped with Insight X3 laser (tuning range 680–1300 nm and fixed 1045 nm) was used for all *in vivo* imaging studies. 920 nm and 1045 nm laser excitation were set for imaging astrocytes and neurons, respectively. Imaging was performed ∼70–200 μm below the dura for astrocytic calcium resonant scanning, ∼100-300 μm below the dura for neuronal calcium imaging at layer 2/3. An Olympus 25×, 1.05 NA water-immersion objective lens with de-gassed ultrasound coupling gel was used for imaging. Emitted light was routed by a 570 nm long-pass dichroic mirror through a 495-540 nm filter (green channel) or a 575-645nm filter (red channel), respectively, and detected with GaAsP-PMTs. For each mouse, three regions of interest (ROIs) were recorded for each condition. Neuronal and astrocytic calcium dynamics were sampled at 15 and 5 Hz for 3min, respectively. Images were acquired at 2× optical zoom and 512×512-pixel resolution. For imaging in anesthetized mice, an air tube was installed in front of the mouse nose. Isoflurane level was maintained at 0.5-1% throughout the imaging session. For imaging in awake animals, mice were habituated to head fixation for at least three days prior to recording.

### 4. Image processing and calcium data analysis

#### 4.1 Preprocessing

Format conversion: The code (oir2tif.m) was developed using MATLAB’s toolbox to convert ‘.oir’ format image files generated by the Olympus multiphoton microscope into ‘.tif’ format.

Motion correction of calcium imaging data: the ‘MotionCorrect’ function from the open-source CaImAn package^42^ available on GitHub (https://github.com/flatironinstitute/CaImAn) was utilized. By adjusting the parameters according to the image resolution and executing the program, the motion-corrected video files can be saved in ‘.tif’ format.

#### 4.2 Calcium data analysis

For astrocytic calcium analysis, the open-source program AQuA2^43,44^, available on GitHub (https://github.com/yu-lab-vt/AQuA2), was used to process astrocytic calcium imaging data. Motion-corrected videos were imported into the program, and parameters were adjusted to apply noise-reduction filtering. AQuA2 identifies regions of activity (ROA) within the images by detecting changes in pixel intensity between frames. It then saves both the average intensity changes and the relative intensity changes (ΔF/F) for each region, enabling characterizing calcium responses. A custom-built MATLAB script (UniqeEvents.m) was then developed to process the ‘.mat’ files generated by AQuA2. This script was designed to identify and remove overlapping or duplicate ROAs. The effective radius R_eff_ of each ROA was approximated based on its calculated area S (*S* = π*R*^2^_*eff*_), and the distance between the geometric centroids of two ROA was compared to the sum of their effective radii to assess overlap. Duplicate regions were removed, and the ΔF/F data for the remaining regions were saved for subsequent statistical analysis.

For neuronal calcium analysis, the neuronal calcium imaging data were imported into the ‘demo_pipeline’ function of the open-source CaImAn package. After adjusting the parameters to suit the video imaging data, the program was executed. The resulting ΔF/F data were saved for subsequent data analysis.

##### Cellular calcium transients analysis

The saved ΔF/F data were consolidated and imported into a custom Python program (dffAnalysis_Findpeaks.py) for analyzing cellular calcium transients. To enhance computational efficiency, the data were downsampled by applying an appropriate step size and window size. Peaks in the downsampled data were identified using the ‘find_peaks’ function, with parameters such as minimum peak height and duration adjusted as needed. This analysis provided the magnitude and timing of calcium response peaks. The identified peaks were then categorized into three phases based on their timing relative to ultrasound stimulation: pre, during, and post-ultrasound stimulation. For each phase, the average peak magnitude, peak frequency, and area under the curve (AUC) of the ΔF/F data were calculated for further analysis. This approach was used to analyze both neuronal and astrocyte calcium dynamics under two different ultrasound stimulation conditions.

##### K-means Clustering Analysis

The AUC data for astrocytes or neurons under different ultrasound stimulation conditions were imported into a custom Python program for clustering analysis (dffAnalysis_Cluster.py). The ‘kmeans.inertia_’ function was used to evaluate clustering performance, and an elbow plot was generated to determine the optimal number of clusters. K-means clustering was then performed on the entire dataset to qualitatively assess calcium signal patterns. The clustered (annotated) data were used for further statistical analysis.

##### Z-score

The ΔF/F data obtained under different ultrasound stimulation conditions were imported into a custom Python program (dffAnalysis_Zscore.py) to calculate the z-score for each cell’s ΔF/F data as follows:

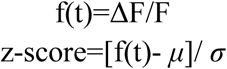

where *μ* is the mean of the ΔF/F data population for each cell, *σ* is the standard deviation of the ΔF/F data population for each cell. Cells were then sorted based on the average intensity during ultrasound stimulation. A z-score heatmap was generated to visualize the sorted z-scores, providing an intuitive representation of the calcium response dynamics under different stimulation conditions over time.

Heatmaps showing the number of calcium responses and spatial variability in different conditions: a custom MATLAB script (HeatMapsInDifferentConditions.m) was developed to process the ‘.mat’ files generated by AQuA2. Based on the ‘.mat’ files, the program identifies regions exhibiting calcium responses and generates heat maps by quantifying the frequency of calcium events across different regions during distinct time periods (Before US, During US, and After US). Additionally, a time-dependent standard deviation projection of the video images is produced to further visualize the spatial variability of calcium activity.

### 5. Evaluation of the laser power output and thermal effects

To estimate the potential thermal effect of two photon imaging and focused ultrasound stimulation, a thermocouple (K-style, connected to an AI-5500 digital thermometer, UDIAN) was implanted 1 mm deep into the S1 cortex of the harvested brain with intact cranial window and head bar to measure the temperature change. A total of 3 min of recording time under two photon imaging, including 1 min of focused ultrasound exposure was performed by utilizing the same ultrasound parameters for *in vivo* stimulation.

The output power of two-photon (2P) laser at 920 nm and 1045 nm excitation wavelength was measured by a laser power meter (Thorlabs PM100D) under the 25x objective. The laser settings were kept the same as the 2P imaging sessions.

## 6. Statistics

Statistical analyses were performed using GraphPad Prism software (version 10.0.3 for MacOS, GraphPad Software, LLC). All the data in this study are represented as mean ± SEM. *N*-values represent individual cells or active events from individual mice. To compare percentages of number and area of active events in astrocytic calcium dynamics, and the average amplitude (ΔF/F), frequency (Hz) and AUC of calcium transients in both neuronal and astrocytic recordings between mice groups, two-way ANOVAs or mixed-effects models were employed, followed by Dunnett’s multiple comparisons test. A significance level of ⍺ = 0.05 was used for all analyses.

## Results

### 1. Experimental System and Characterization

To enable concurrent ultrasound stimulation and optical imaging of astrocytic and neuronal calcium dynamics, we integrated a low-frequency focused ultrasound (FUS) transducer with a two-photon (2P) laser scanning fluorescence microscope (Figure 1A and Figure S1). We randomly employed two sets of ultrasound parameters with different intensities and low duty cycles and tested if similar or different responses in astrocytes and neurons could be elicited. The FUS was delivered at a foundamental frequency of 0.521MHz and spatial-peak pulse-average intensity (I_SPPA_) of 0.91 or 1.5 W/cm^2^. A 4 mm cranial window, sealed with dental cement and a glass coverslip, provided clear optical access for 2P imaging (Figure 1A). To verify the FUS stimulation profile, we positioned a hydrophone beneath a glass coverslip in a water tank. The measured pressure maps in the lateral (X-Y) and vertical (X-Z) planes indicated a focused acoustic field about 2 mm in diameter for −6 dB acoustic pressure and 2 mm in focal depth (Figure 1B). To mitigate the heating effect of FUS, a low duty cycle of 5% or 10% was used in our experiments. We also measured the temperature change in the absence and presence of FUS under different stimulation parameters. A temperature probe implanted in the mouse cortex revealed that 1min of FUS at intensities of 0.91 or 1.5 W/cm^2^ did not induce temperature increase relative to the group without FUS stimulation (Figure 1C-D). Notably, 2P imaging itself at 920 nm or 1045 nm excitation produced transient cortical temperature rise of approximately 0.4 ℃ and 0.6 ℃, respectively. These increases returned to baseline promptly after the imaging session ended (Figure 1C-D). The data confirms that the temporary temperature increase in mouse cortex was associated with extended 2P laser excitation, and independent of FUS stimulation used in our study.

**Figure 1:**
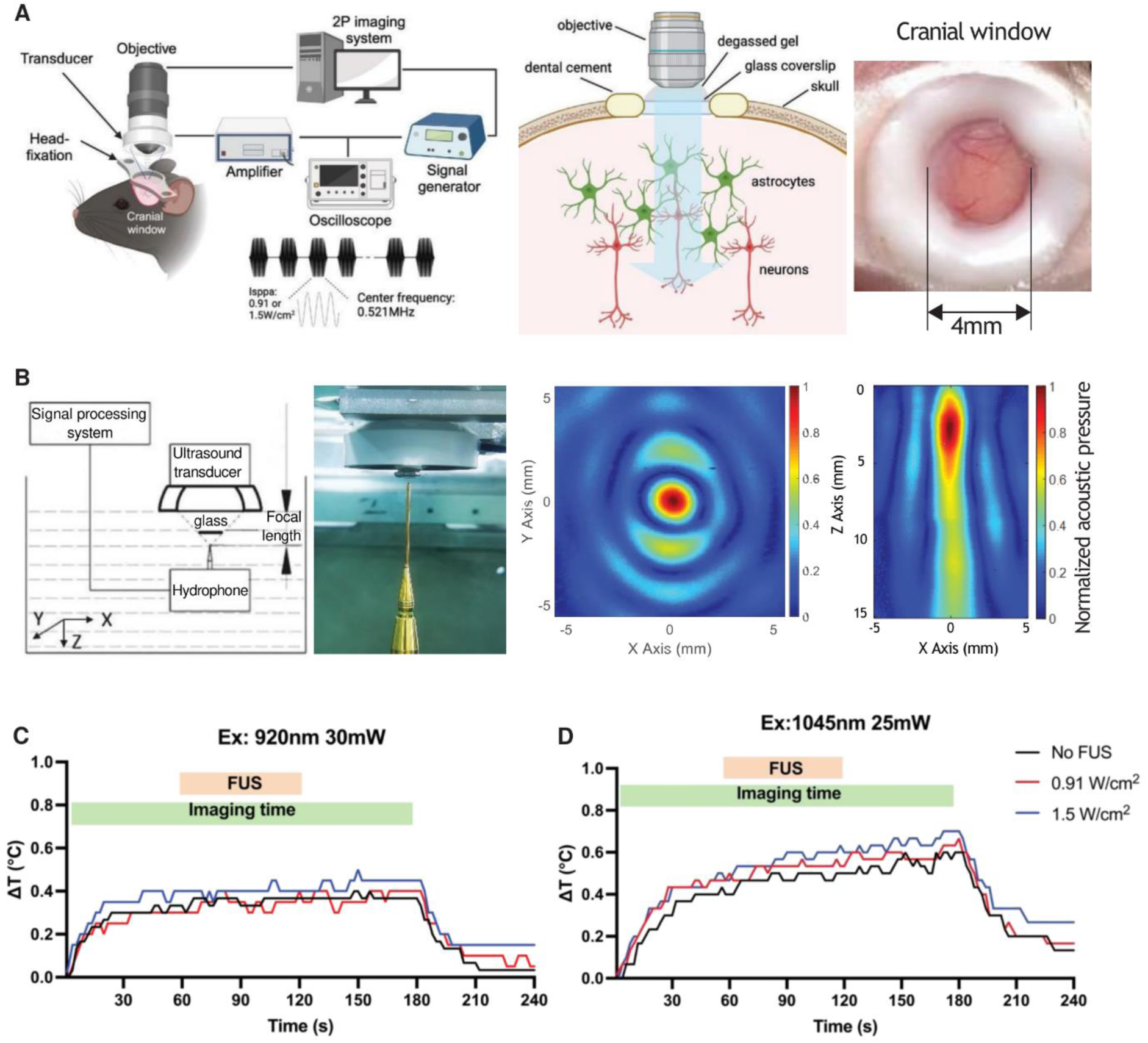
Experimental system integrating FUS stimulation and 2P imaging and its characterization. (A) Schematic of experimental setup for simultaneous *in vivo* 2P imaging and focused ultrasound stimulation. A transducer was coupled to a cranial window (4 mm diameter) via a layer of degassed ultrasound gel, enabling ultrasound stimulation to the S1 cortex layer 2/3 while maintaining optical access for imaging astrocytes and neurons. FUS was delivered at a fundamental frequency of 0.521MHz with an intensity I_SPPA_ of 0.91 or 1.5 W/cm^2^. (B) Characterization of the acoustic field of the customized ultrasound transducer through the glass coverslip using a hydrophone in degassed water tank. The normalized acoustic pressure distribution in the XY and XZ focal planes were shown in the right two panels. (C) Temperature changes (ΔT) of the mouse S1 cortex in response to FUS (orange bar) at 0.91 or 1.5 W/cm² and 2P laser excitation at wavelength of 920 nm, power of 30 mW and 5 Hz imaging frame rate (settings used for astrocytes). (D) Temperature changes (ΔT) of the mouse S1 cortex in response to FUS (orange bar) at 0.91 or 1.5 W/cm² and 2P laser excitation at wavelength of 1045 nm, power of 25 mW and 15 Hz imaging frame rate (settings used for neurons). Green bars indicate imaging time.

### 2. FUS promoted astrocyte calcium activity in awake mice

To investigate how FUS influences the calcium signaling of astrocytes *in vivo*, the mice were stereotaxic injected with the recombinant adeno-associated virus encoding GCaMP6f in astrocytes into layer 2/3 of the somatosensory cortex (S1). We were able to detect spontaneous calcium activity in astrocyte microdomains by 2P imaging in awake mice (Figure 2A). Individual astrocyte microdomain calcium tracing and temporal projection of astrocyte calcium dynamics in the imaging field showed increased astrocyte calcium responses during FUS stimulation at both intensities of 0.91 and 1.5 W/cm^2^ (Figure 2A-B). FUS stimulation at I_SPPA_ of 1.5 W/cm^2^ elicited higher average calcium peaks compared to the other group (Figure 2C). Quantification of the spatial ratio and normalized number of regions of activity showed no significant change during, and after FUS compared to the baseline before FUS among the control and FUS-treated awake mice (Figure 2D-E). Indeed, significant increases of amplitude, frequency, and area under the curve of calcium transients (temporal integral) in astrocyte subregions were observed during FUS stimulation compared to those before FUS application (Figure 2F-H). We also found elevated calcium response in the control group during recording, most likely due to the accumulated heating effect of 2P laser at 920 nm excitation wavelength (Figure 2F-H). Further analysis for the calcium responses in individual astrocyte microdomain revealed that FUS stimulation at 1.5 W/cm^2^ elicited the strongest activation effect (Figure 3A, Figure S3). Specifically, FUS stimulation at 1.5 W/cm^2^ further promoted the calcium response in astrocytes with 39.5% ROA population activated, compared to 25.6% in No FUS and 31.7% in FUS at 0.91 W/cm^2^ group (Figure 3B-C). Collectively, these data indicate that FUS promotes robust increase in astrocyte calcium activity in awake mice.

**Figure 2:**
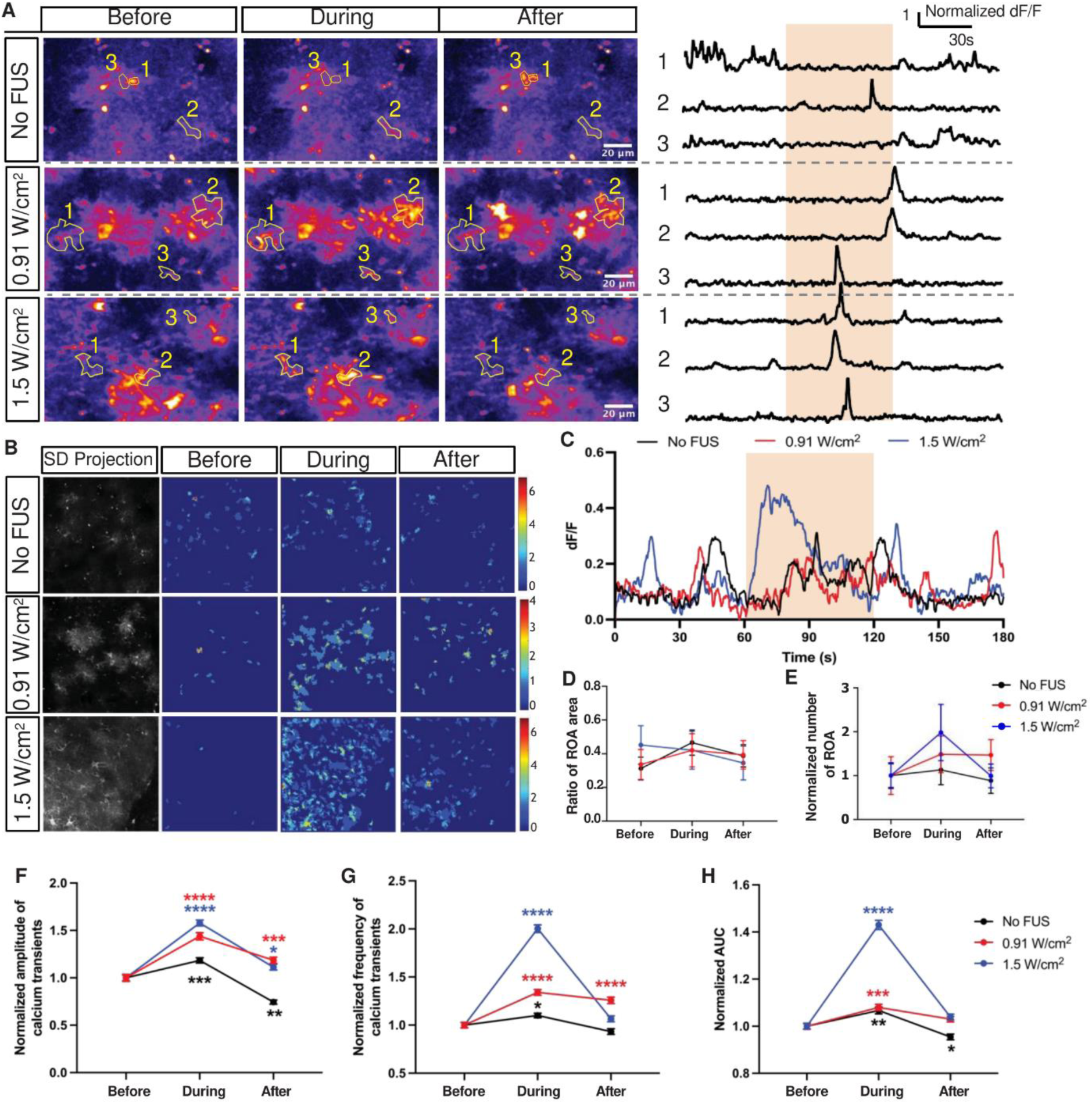
*in vivo* 2P imaging of astrocyte calcium dynamics in response to FUS in awake mice. (A) Exemplary standard deviation projection of 2P astrocyte calcium imaging and normalized individual dF/F traces before, during, and after FUS stimulation at intensities of 0 (No FUS), 0.91, or 1.5 W/cm^2^ in awake mice, respectively. A hotter color depicts larger change. Yellow outlines marked by 1, 2 and 3 indicate individual astrocyte subregions. The orange box on the right depicts the timing of ultrasound stimulation for 1 min. (B) Representative heatmaps of astrocyte calcium dynamics in the imaging field before, during, and after FUS stimulation at intensities of 0 (No FUS), 0.91, or 1.5 W/cm^2^, respectively. Left grayscale images were the standard deviation temporal projection of astrocyte calcium signals. The color bars indicate the number of calcium peaks. (C) Average astrocyte calcium response over time from all awake mice under FUS stimulation at intensities of 0 (No FUS), 1.5, or 0.91 W/cm^2^, respectively. (D) The ratio of ROA area to the total events area, where the event area quantification is detailed in Figure S2. (E) the normalized change of the number of astrocyte ROA from awake mice before, during, and after FUS stimulation at intensities of 0 (No FUS), 0.9, or 1.5 W/cm^2^, respectively. (F) Normalized amplitude of calcium transients, (G) normalized frequency of calcium transients, and (H) normalized area under the curve (AUC) of astrocyte calcium responses before, during and after FUS stimulation at intensities of 0 (No FUS), 0.91, or 1.5 W/cm^2^, respectively. Data are presented as mean±SEM. *n =* 1567 (No FUS), 1377 (0.91 W/cm^2^) and 1636 (1.5 W/cm^2^) astrocyte responsive events from 7 awake mice. Significant differences were determined by two-way ANOVA with Tukey’s and Dunnett’s multiple comparisons tests. \**p*<0.05, \*\**p*<0.01, \*\*\**p*<0.001, \*\*\*\**p*<0.0001.

**Figure 3:**
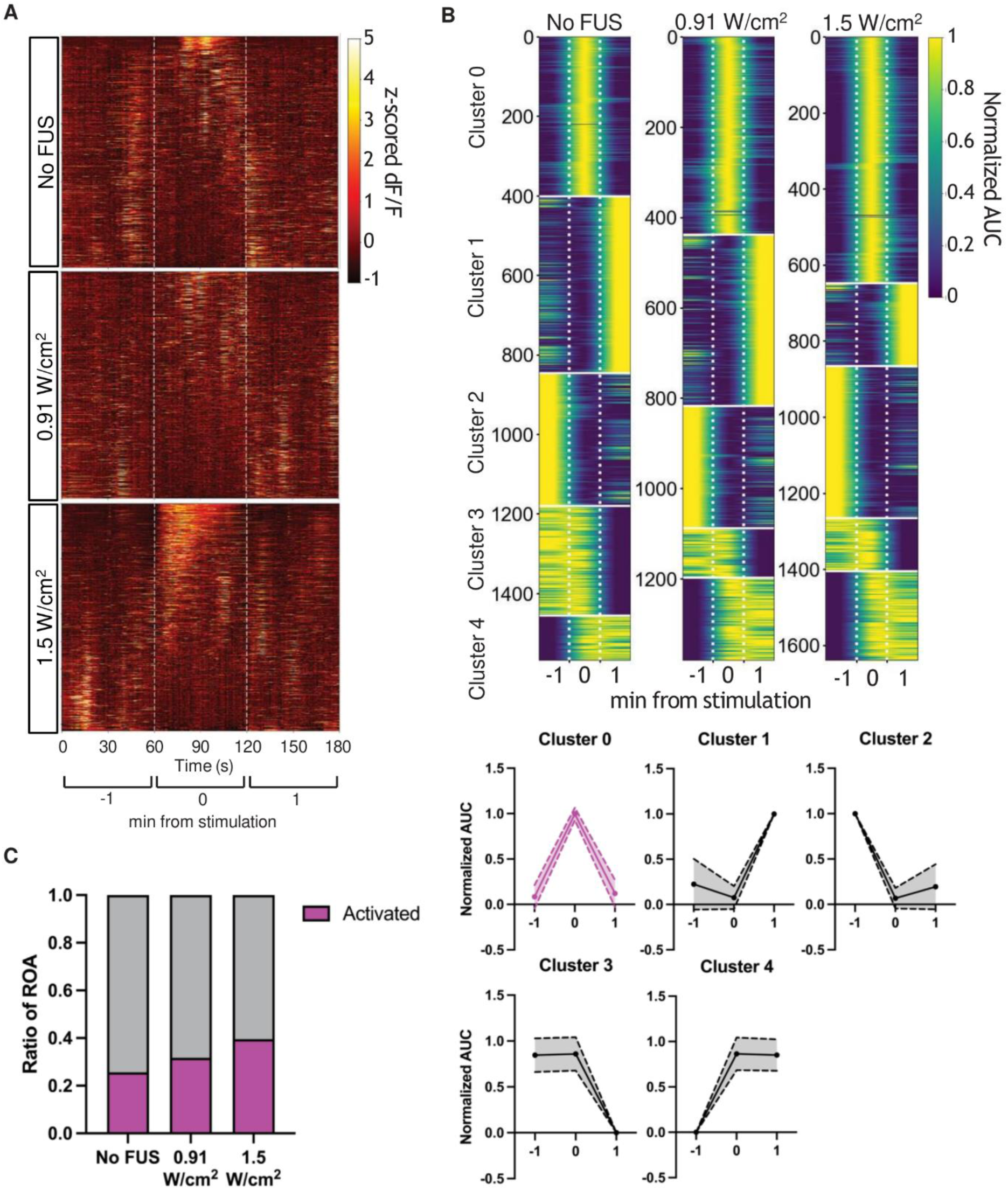
Population analysis of astrocyte calcium dynamics in response to FUS in awake mice. (A) Heatmaps of the normalized z-scored dF/F of astrocyte calcium dynamics over time under three conditions: No FUS, 0.91 W/cm², and 1.5 W/cm² FUS. Each row represents an individual astrocyte ROA responsive calcium event. The white dashed lines indicates the time window for FUS stimulation. (B) Clustering analysis of astrocytic calcium responses. Upper panels show grouped activity traces based on K-means clustering, categorized into five distinct response clusters (Cluster 0–4) across FUS conditions. Heatmaps represent individual astrocyte ROA calcium active events (y-axis) over time (x-axis) for each cluster. Bottom panels display mean normalized calcium response AUC (before, during and after FUS) for each cluster, representing distinct temporal response profiles. Cluster 0 (magenta) exhibits transient activation during FUS stimulation (0 min). Solid lines depict the mean value while the shadowed regions represent SD. (C) The ratio of activated astrocyte ROA calcium events (magenta, Cluster 0) to total responsive astrocyte events (grey) under no FUS, 0.91 W/cm², and 1.5 W/cm² FUS.

### 3. FUS-activated astrocyte calcium responses were less pronounced in lightly anesthetized mice

Calcium levels of astrocyte change according to the sleep/wakefulness states and associate with cortical state switching^45–47^. FUS has been shown to induce different neural activities depending on the behavioral states of the animal^31,48^. However, whether FUS stimulation modulates astrocyte calcium dynamics in anesthetized animals, similar to or different from the awake mice, remain unknown. Previous studies have found that general anesthesia depressed calcium responses in astrocytes and triggered activity-related gene expression changes^49–51^. To determine if FUS could promote astrocyte calcium activity in low-activity environment, we next examined the astrocytic calcium dynamics before, during, and after FUS stimulation in isoflurane anesthetized mice (Figure 4A). The mice were first induced to anesthesia in 3% isoflurane and then maintained in the lightly anesthetized state at 0.5% isoflurane. Temporal projection and tracing of astrocyte microdomains showed no obvious change of overall astrocyte calcium responses in FUS groups (Figure 4B-C, Figure 5A). Consistent with awake data, we observed no significant change in either the total area or the number of ROA in astrocytes from lightly anesthetized mice compared to baseline value (Figure 4D-E). Despite unchanged overall spatial activation, FUS stimulation at 0.91 and 1.5 W/cm^2^ produced a robust increase in the amplitude of astrocyte calcium transients (Figure 4F). These effects persisted after FUS cessation, leading to significant increases in both event frequency and area under the curve (AUC) of the calcium transients (Figure 4G-H). However, the frequency and AUC of astrocyte microdomain calcium transients also increased towards the end of imaging in the absence of FUS stimulation (Figure 4G-H), though to a smaller extent compared to that with FUS stimulation (Figure S4), indicating heating generated by 2P laser excitation and imaging itself could confound the astrocyte calcium dynamics in lightly anesthetized mice. In our study, care has been taken to reduce the heat effects by employing a slower framing rate of 5 Hz at this condition. In addition, clustering analysis showed that FUS stimulation only raised the percentage of activated ROA population by 4.6–4.8%, comparing to the 23.3% of spontaneously activated ROA in astrocytes in the absence of FUS stimulation (Figure 5B-C), less than the awake state. These data suggest that FUS could still promote astrocyte calcium responses in anesthetized mice (Figure S4), and the effect is less pronounced than in the awake state.

**Figure 4:**
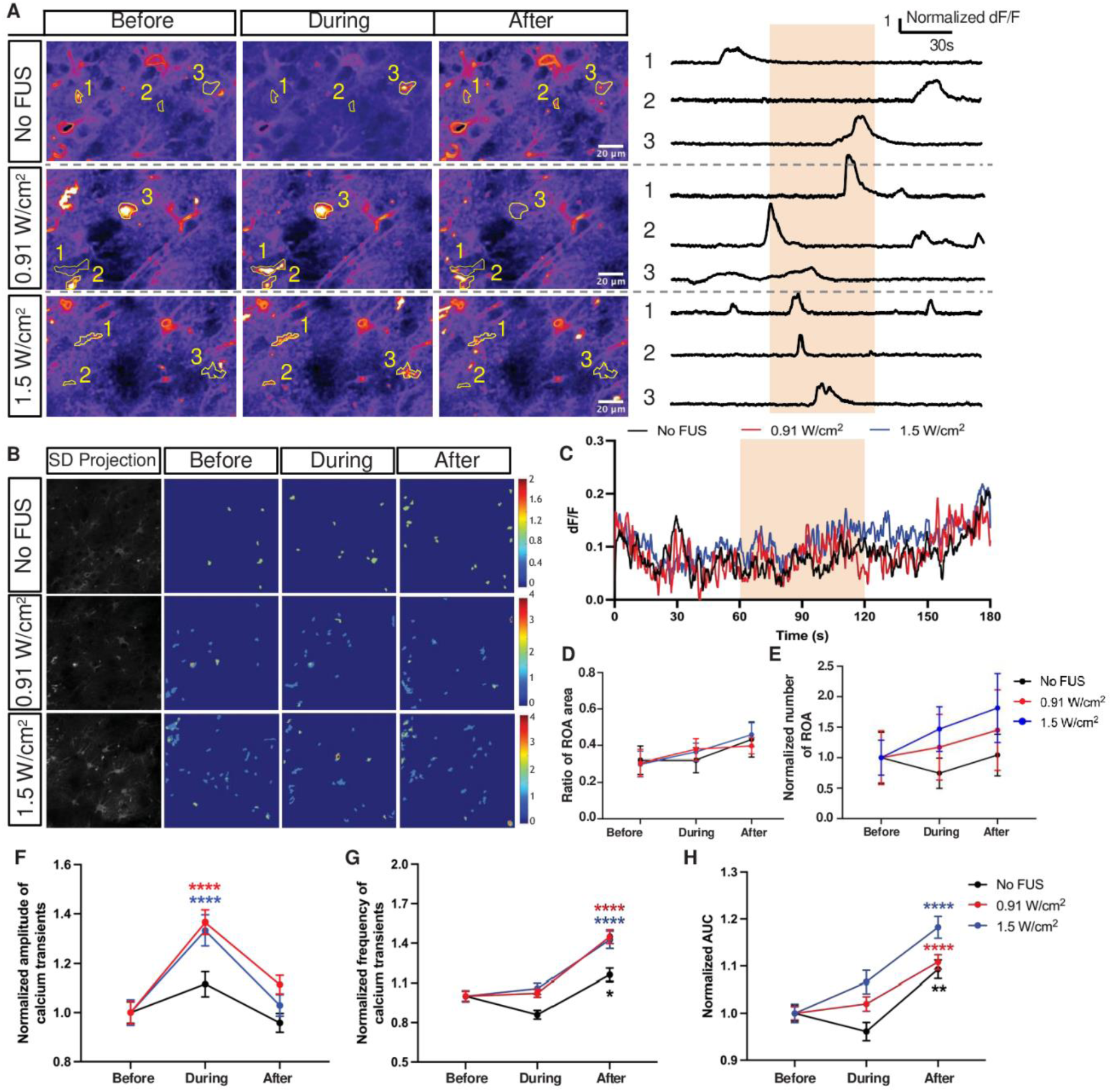
*in vivo* 2P imaging of astrocyte calcium dynamics in response to FUS in lightly anesthetized mice. (A) Exemplary pseudocolor images of the standard deviation projection of astrocyte calcium fluorescent with 2P imaging and normalized individual dF/F traces before, during (yellow shaded region), and after FUS stimulation at intensities of 0 (No FUS), 0.91, or 1.5 W/cm^2^ in lightly anesthetized mice, respectively. A hotter color depicts larger change. Yellow outlines indicate individual astrocyte subregions. (B) Representative heatmaps of astrocyte calcium dynamics in the imaging field before (−1min), during (0min), and after (+1min) FUS stimulation at intensities of 0 (No FUS), 0.91, or 1.5 W/cm^2^, respectively. Left grayscale images were the standard deviation temporal projection of astrocyte calcium signals. The color bars indicate the total number of calcium peaks. (C) Average astrocyte calcium responses over time from light anesthetized mice under FUS stimulation at intensities of 0 (No FUS), 0.91, or 1.5 W/cm^2^, respectively. Orange box indicates the time of FUS stimulation. (D) The ratio of ROA area to the total events area, where the event area quantification is detailed in Figure S2. (E) the normalized change of the number of astrocyte ROA from lightly anesthetized mice before, during, and after FUS stimulation at intensities of 0 (No FUS), 0.9, or 1.5 W/cm^2^, respectively. (F) Normalized amplitude of calcium transients, (G) normalized frequency of calcium transients, and (H) normalized area under the curve (AUC) of astrocyte calcium responses before, during, and after FUS stimulation at intensities of 0 (No FUS), 0.91, or 1.5 W/cm^2^, respectively. Data are presented as mean ± SEM. *n* = 691 (no FUS), 904 (0.91 W/cm^2^) and 563 (1.5 W/cm^2^) astrocyte responsive events from 5 lightly anesthetized mice. Significant differences were determined by two-way ANOVA with Tukey’s and Dunnett’s multiple comparisons tests. *p<0.05, **p<0.01, ***p<0.001, ****p<0.0001.

**Figure 5:**
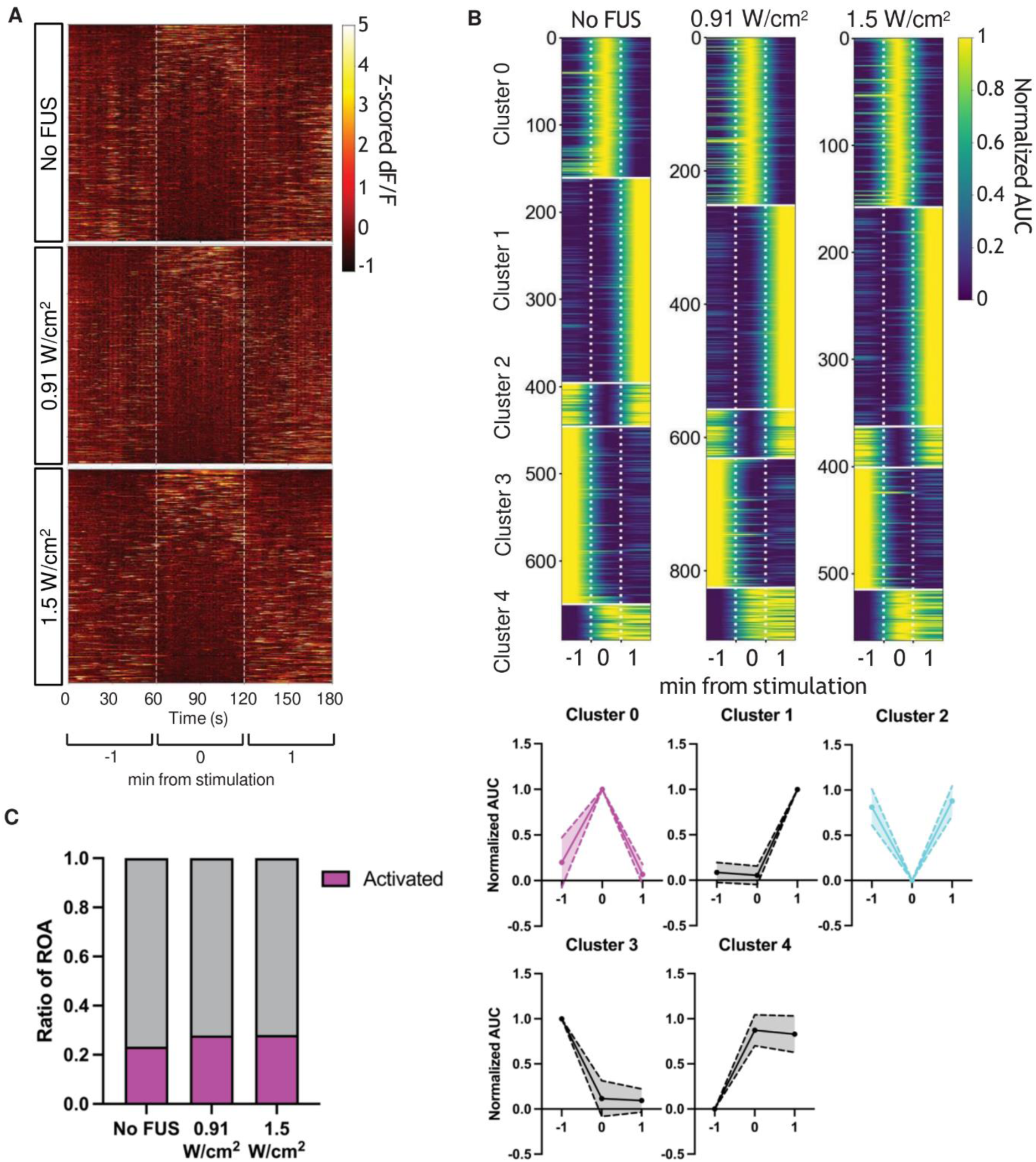
Population analysis of astrocyte calcium dynamics in response to FUS in lightly anesthetized mice. (A) Heatmaps of the normalized z-scored dF/F of astrocyte calcium dynamics over time under three conditions: No FUS, 0.91, and 1.5 W/cm² FUS. Each row represents an individual astrocyte ROA responsive calcium event. (B) Clustering analysis of astrocytic calcium responses in lightly anesthetized mice. Upper panels show grouped activity traces based on K-means clustering, categorized into five distinct response clusters (Cluster 0–4) across FUS conditions. Heatmaps represent individual astrocyte ROA calcium active events (y-axis) over time (x-axis) for each cluster. Bottom panels display mean normalized calcium response AUC for each cluster, representing distinct temporal response profiles. Cluster 0 (magenta) exhibits transient activation during FUS stimulation. Solid lines depict the mean value while the shadowed regions represent SD. (C) The ratio of activated astrocyte ROA calcium events (magenta, Cluster 0) to total responsive astrocyte events (grey) under No FUS, 0.91 W/cm² and 1.5 W/cm² FUS in lightly anesthetized mice.

### 4. FUS suppressed neuronal calcium activity in awake mice

Previous findings show that astrocyte calcium dynamics could shape neuronal network activity. Our study has demonstrated that FUS promoted astrocyte calcium dynamics. Next, we investigate the relevant neuronal activity change *in vivo* at the same FUS parameters. To examine the neuronal calcium dynamics before, during, and after FUS stimulation *in vivo*, mice were stereotaxic injected with the recombinant virus expressing jRGECO1a in layer 2/3 neurons in S1 cortex for calcium imaging (Figure 6A). Calcium dynamics from individual neurons and the imaging filed were analyzed (Figure 6B-C). Average calcium traces showed no significant difference among the groups (Figure 6D). The automatic calcium signal analysis pipeline was able to detect around 80% of neuronal calcium responses across groups (Figure 6E). Although the amplitude of neuronal calcium transient increased after FUS stimulation at 0.91 W/cm^2^ compared to the neuronal calcium transients before FUS stimulation (Figure 6F), the frequency of calcium transient significantly decreased during FUS at both 0.91 and 1.5 W/cm^2^ and recovered afterwards (Figure 6G). Moreover, both FUS paradigms significantly suppressed neuronal calcium activity as indicated by area under the curve (Figure 6H). Moreover, we investigated the activity distribution of neuronal calcium transients to determine which functional subpopulation was mostly affected by ultrasound stimulation (Figure 7A-B). Five activity clusters were identified from 1600-1700 neurons recorded from ten mice for each group by K-means clustering analysis (Figure 7B). Based on the changes of normalized AUC, we were able to identify activated (cluster 3) and inhibited (cluster 4) functional subpopulation of responsive neurons. Notably, FUS stimulation at 0.91 W/cm^2^ significantly reduced the activated neuronal functional subpopulation (Figure 7B-C and Figure S5), supporting the previous finding of FUS-dependent suppression of neuronal calcium response.

**Figure 6:**
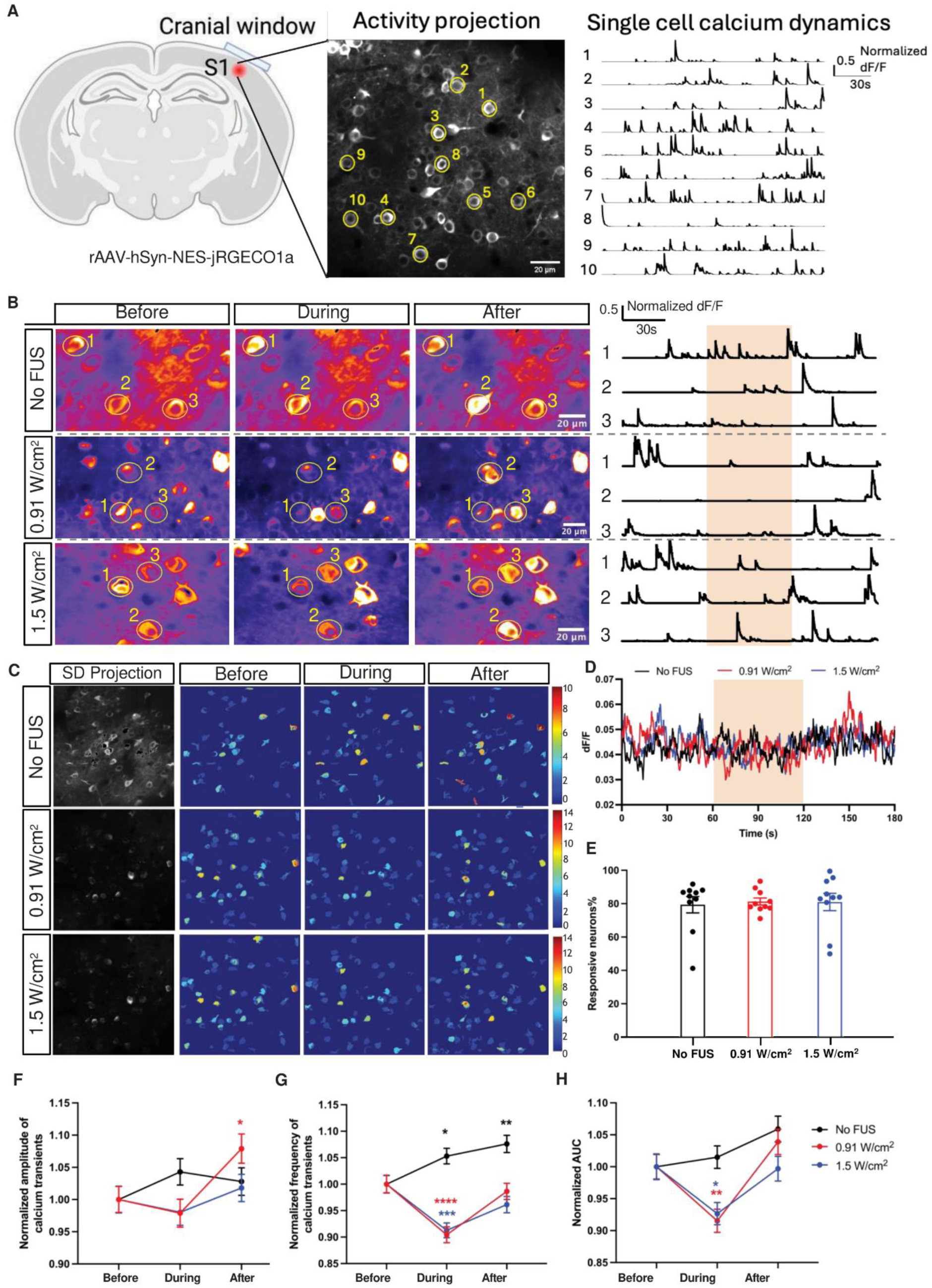
*in vivo* 2P imaging of neuronal calcium dynamics in response to FUS in awake mice. (A) Representative 2P neuronal calcium imaging of mouse S1 cortex. Right panel shows exemplary calcium transient traces of the neurons labeled with yellow circles in the middle panel. (B) Exemplary pseudocolor images of the standard deviation projection of neuronal calcium response with 2P imaging and normalized individual dF/F traces before, during, and after FUS stimulation at intensities of 0 (No FUS), 0.91, or 1.5 W/cm^2^ in awake mice, respectively. A hotter color indicates larger variation. The orange box over the traces indicates the time of FUS stimulation. (C) Representative heatmaps of calcium dynamics in the imaging field before, during, and after FUS stimulation at intensities of 0 (No FUS), 0.91, or 1.5 W/cm^2^, respectively. Left grayscale images were the standard deviation temporal projection of neuronal calcium signals. The color bars on the right indicate the total number of calcium peaks. (D) Average neuronal calcium responses over time from all awake mice under FUS stimulation at intensities of 0 (No FUS), 0.91, or 1.5 W/cm^2^, respectively. (E) Percentage of responsive neurons in awake mice under each condition. (F) Normalized amplitude of calcium transients, (G) normalized frequency of calcium transients, and (H) normalized area under the curve (AUC) of calcium responses before, during, and after FUS stimulation at intensities of 0 (No FUS), 0.91, or 1.5 W/cm^2^, respectively. Data are presented as mean ± SEM. *n* = 1671 (No FUS), 1650 (0.91 W/cm^2^) and 1611 (1.5 W/cm^2^) neurons from 10 mice. Significant differences were determined by two-way ANOVA with Tukey’s and Dunnett’s multiple comparisons tests. \**p*<0.05, \*\**p*<0.01, \*\*\**p*<0.001, \*\*\*\**p*<0.0001.

**Figure 7:**
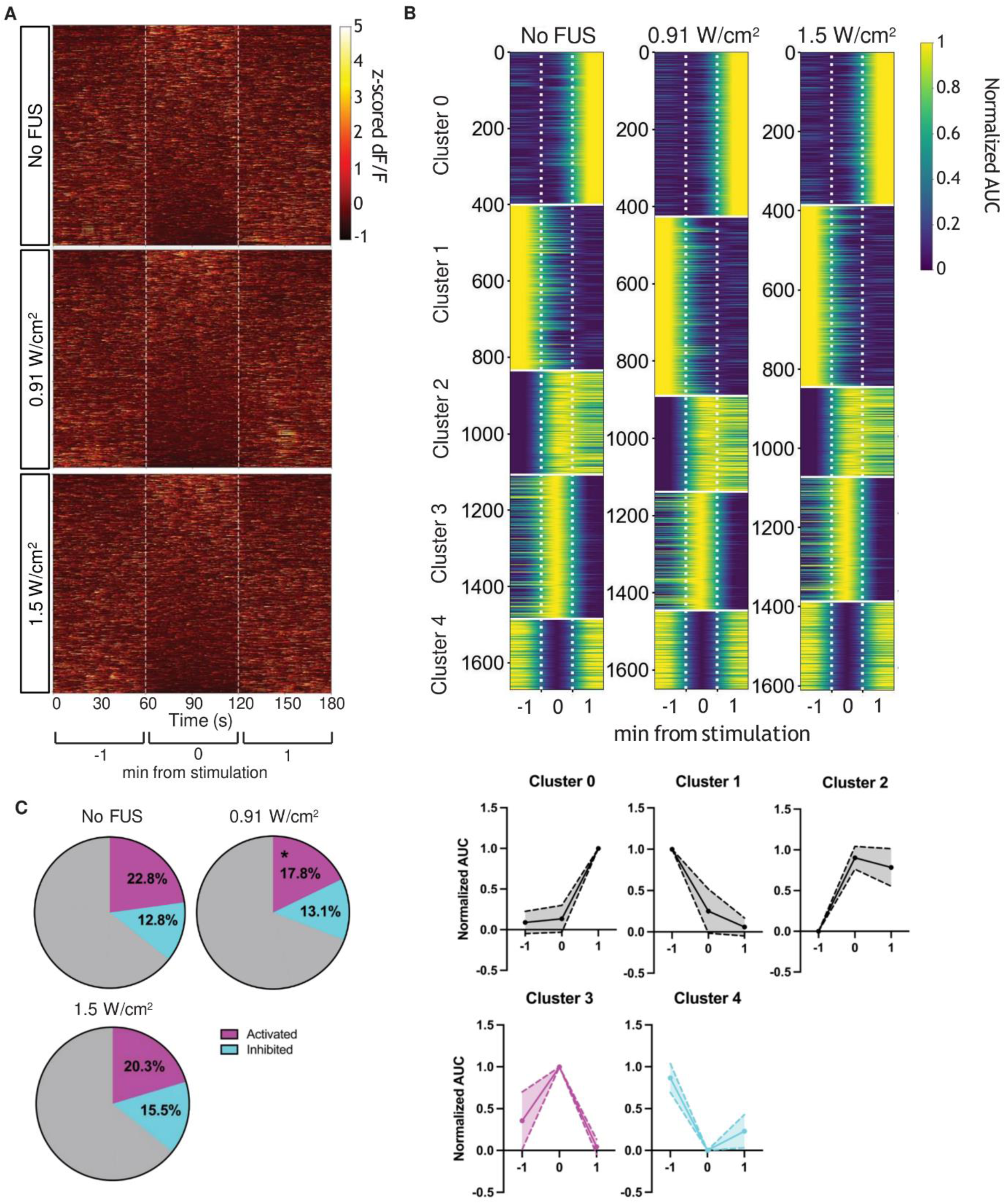
Population analysis of neuronal calcium dynamics in response to FUS in awake mice. (A) Heatmaps of the normalized z-scored dF/F of neuronal calcium dynamics over time in response to No FUS, FUS of 0.91 W/cm², and 1.5 W/cm² conditions. Each row represents an individual neuron, showing temporal response patterns before (−1 min), during (0 min), and after (+1 min) FUS stimulation. (B) Clustering analysis of neuronal calcium responses in awake mice. Upper panels show grouped activity traces based on K-means clustering, categorized into five distinct neuronal response clusters (Cluster 0–4) across FUS conditions. Heatmaps show the normalized AUC values for each cluster (x-axis: time, y-axis: individual neurons) under No FUS, 0.91 W/cm², and 1.5 W/cm² FUS. The average response profiles (solid lines) with SD labeled by shadow area for each cluster are shown in the bottom panel. The cluster 3 (magenta) and cluster 4 (cyan) are identified as activated and inhibited neuronal calcium response during FUS, respectively. (C) The proportions of activated (magenta, cluster 3), inhibited (cyan, cluster 4), and other responsive (grey) neurons under No FUS, 0.91 W/cm², and 1.5 W/cm² FUS. Data are presented as the means of each group in the pie chart. *n* = 10 mice for each condition. Significant differences were determined by two-way ANOVA with Dunnett’s multiple comparisons test. *p<0.05.

### 5. Neuronal calcium dynamics were not significantly perturbed by FUS in anesthetized mice

To determine if FUS could alter neuronal calcium dynamics in mice with inherently suppressed neuronal activity and calcium transient, we recorded the jRGECO1a fluorescent signals in mice under light anesthesia with isoflurane (Figure 8A-C). Only 20-25% neurons in lightly anesthetized mice were detected to be responsive via the automatic pipeline, which is significantly less than neurons in awake mice (Figure 8D). Compared to the control group, no obvious change in neuronal calcium response was found in FUS treated mice (Figure 8E, Figure 9A). The average frequencies of neuronal calcium transients in mice decreased upon FUS stimulation at both 0.91 W/cm^2^ and 1.5 W/cm^2^ compared to the baseline frequency prior to FUS stimulation (Figure 8F). There are no significant changes in the area under the curve of neuronal calcium response to FUS stimulation in anesthetized mice (Figure 8G). We next performed the functional clustering analysis from the 600-700 responsive neurons detected in eleven mice (Figure 9B). The distribution of neuronal calcium activity subpopulations in FUS-treated mice is similar to the control group, without significant changes in activated or inhibited neuronal subpopulations (Figure 9B-C and Figure S6). Taken together, these results show that the effect of ultrasound on neuronal calcium activity is insignificant in mice under anesthesia.

**Figure 8:**
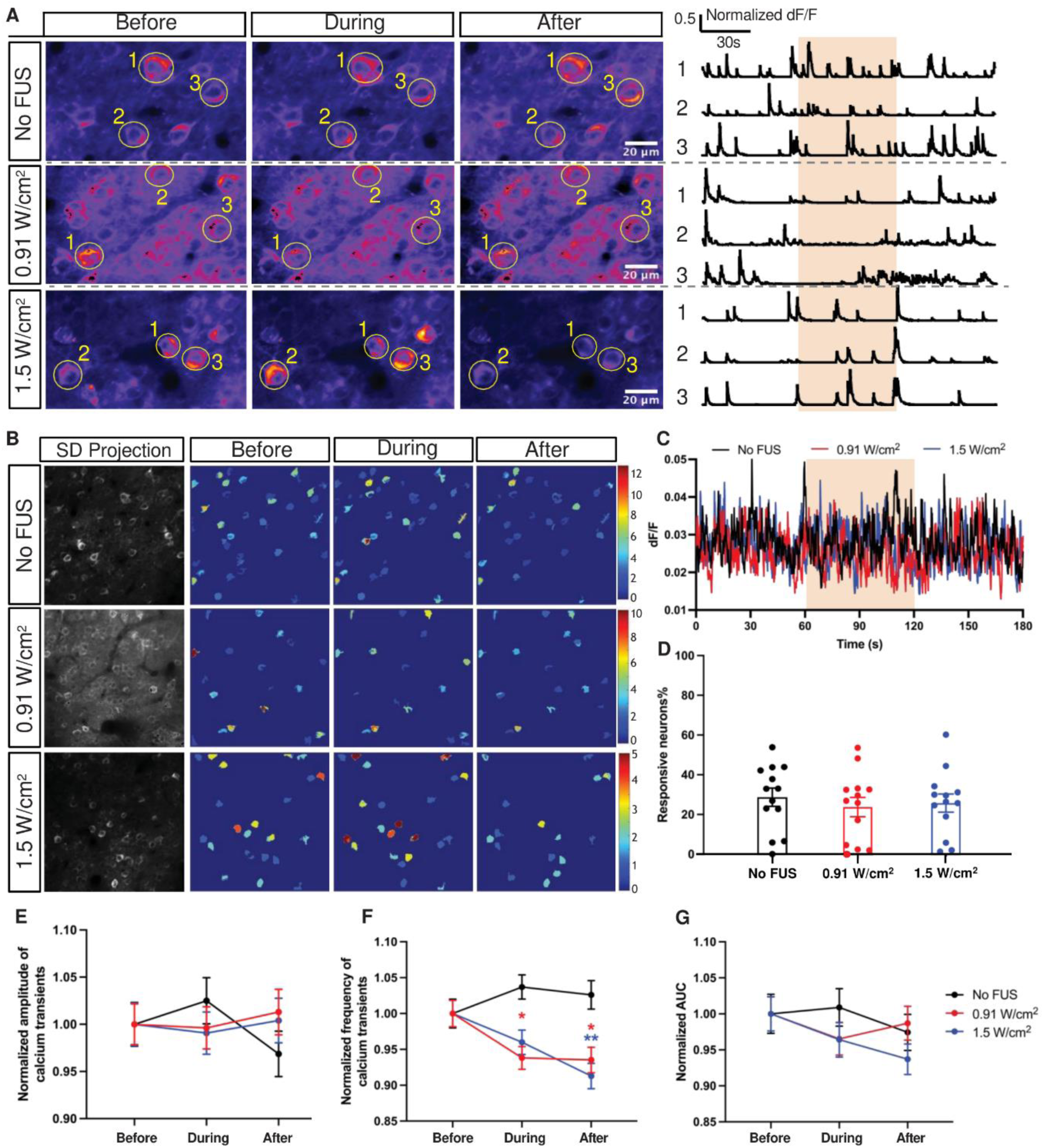
*in vivo* 2P imaging of neuronal calcium dynamics in response to FUS in lightly anesthetized mice. (A) Exemplary standard deviation projection of neuronal calcium fluorescence intensity with 2P imaging and normalized individual dF/F traces for the neurons marked by 1, 2 and 3 in each condition before, during, and after FUS stimulation at intensities of 0 (No FUS), 0.91, or 1.5 W/cm^2^ FUS in lightly anesthetized mice, respectively. A hotter color indicates larger variation. The orange box over the traces indicates the time of FUS stimulation. (B) Representative heatmaps of calcium dynamics in the imaging field in lightly anesthetized mice before, during, and after FUS stimulation at intensities of 0 (No FUS), 0.91, or 1.5 W/cm^2^, respectively. Left grayscale images were the standard deviation temporal projection of neuronal calcium signals. The color bars indicate the number of calcium peaks. (C) Average neuronal calcium response over time from all lightly anesthetized mice under FUS stimulation at intensities of 0 (No FUS), \ 0.91, and 1.5 W/cm^2^, respectively. (D) Percentage of responsive neurons detected by automatic pipelines. Each datapoint represent a mouse. (E) Normalized amplitude of calcium transients, (F) normalized frequency of calcium transients, and (G) normalized area under the curve (AUC) of neuronal calcium responses in lightly anesthetized mice before, during, and after FUS stimulation at intensities of 0 (No FUS), 0.91, or 1.5 W/cm^2^, respectively. Data are presented as mean ± SEM. *n =* 681 (No FUS), 631 (0.91 W/cm^2^) and 653 (1.5 W/cm^2^) neurons from 13 mice. Significant differences were determined by two-way ANOVA with Tukey’s and Dunnett’s multiple comparisons tests. \**p*<0.05, \*\**p*<0.01, \*\*\**p*<0.001, \*\*\*\**p*<0.0001.

**Figure 9:**
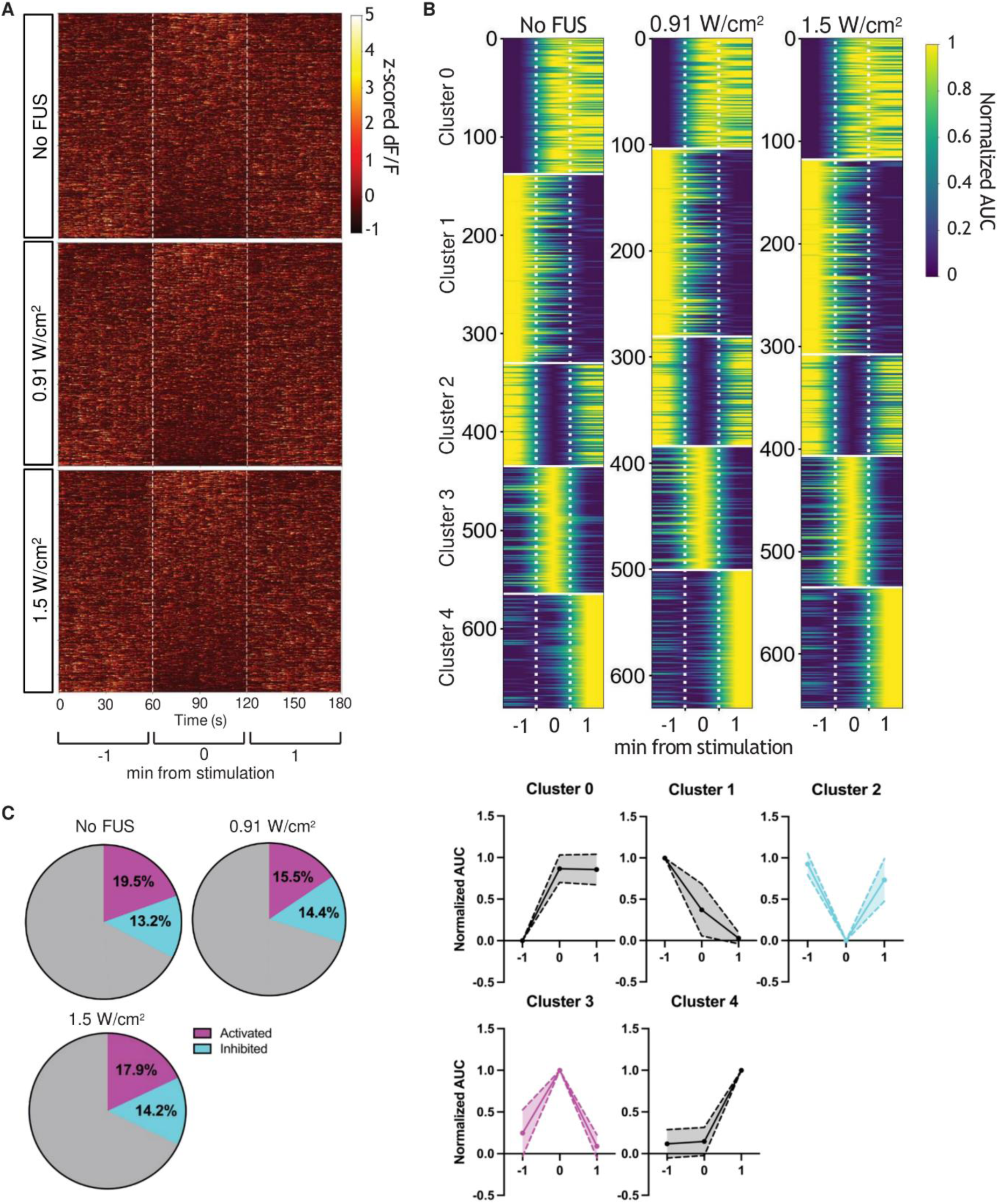
Population analysis of neuronal calcium dynamics in response to FUS in lightly anesthetized mice. (A) Z-scored normalized neuronal calcium activity in lightly anesthetized mice without (No FUS) and with ultrasound stimulation at 0.91 and 1.5 W/cm^2^. The white dashed lines indicate the time window for FUS stimulation. (B) Upper: K-means clustering analysis of neuronal calcium activity in lightly anesthetized mice untreated and treated with FUS stimulation at I_SPPA_ of 0.91 and 1.5 W/cm^2^. The representative changes of normalized AUC values for each cluster are shown in the bottom panel. Cluster 3 (magenta) and cluster 2 (cyan) are identified as activated and inhibited neuronal calcium response, respectively. (C) Ratio of neurons for each functional subpopulation of calcium activity in anesthetized mice untreated and treated with ultrasound stimulation at 0.91 and 1.5 W/cm^2^. Data in the pie charts are presented as Mean, *n* = 13 mice for each condition. Significant differences were determined by two-way ANOVA with Dunnett’s multiple comparisons test.

In summary, our dual-modality system combining low-frequency FUS with two-photon imaging revealed distinct, state-dependent effects on astrocyte and neuronal calcium activity. In awake mice, FUS significantly enhanced the amplitude, frequency, and temporal integrated calcium response of astrocyte microdomains, while suppressing neuronal calcium activity and reducing the proportion of activated neuronal subpopulations. Under light anesthesia, FUS still promoted astrocyte calcium responses but to a lesser extent than in the awake condition, and it had little effect on neuronal calcium dynamics. Collectively, these findings demonstrate that FUS can selectively modulate cortical astrocyte and neuronal calcium activity in a manner dependent on the animal’s arousal state.

## Discussion

Compared with the relatively well-studied effects of ultrasound on neuronal excitability^52–54^, much less is known about how FUS influences astrocyte calcium dynamics. In this study, we investigated the effects of FUS on astrocyte and neuronal calcium dynamics *in vivo*. Our data showed significant increases in the amplitude, frequency, and area under the curve of astrocyte calcium transients during or after FUS stimulation, consistent with previous *in vitro* findings^19,55^. Functional subpopulation analysis revealed an elevated proportion of activated astrocytic ROAs in FUS-treated groups. *In vitro* and *in situ* studies showed that ultrasound could potentially elevate astrocyte calcium levels via mechanosensitive channels, promoting the release of neurotransmitters that modulate adjacent neurons^55–58^. Although our present work did not dissect specific potential mechanosensitive pathways^59–61^, the pronounced astrocyte activation we observed *in vivo* suggests that FUS can potently engage non-neuronal cells in the cortical microenvironment.

Interestingly, while FUS robustly enhanced astrocyte calcium activity in awake mice, its effect in lightly anesthetized mice was present but attenuated. Because isoflurane suppresses synaptic activity and cortical excitability^62,63^, this lower baseline activity may reduce the magnitude of FUS-mediated astrocyte calcium responses. Moreover, previous study showed that general anesthesia suppressed and desynchronized spontaneous calcium transients, as well as sensory-evoked calcium responses in cortical astrocytes^49^. Therefore, the presence of isoflurane may attenuate the FUS-promoted calcium activity in astrocytes as shown in our results.

In contrast to the result for astrocytes, our study showed that neuronal calcium activity in awake mice was selectively inhibited by FUS, primarily at the 0.91 W/cm^2^ intensity. Like astrocytes, this effect diminished in lightly anesthetized mice, suggesting that the baseline suppression from isoflurane could mask FUS-mediated neuronal modulation. Functional clustering analysis revealed that FUS delivered at intensity of 0.91 W/cm^2^ preferentially inhibited intrinsically activated neuronal subpopulations in awake mice. Despite the difficulty of dissociating direct FUS-induced neuronal responses from indirect spontaneous network responses *in vivo*, it was reported that excitatory and inhibitory neurons in anesthetized rodent brains were intrinsically different in response to ultrasound PRF^54^. Further study showed that FUS-mediated effect is likely more heterogenous between individual cells than canonical cell types^64^. It is worth noting that we recorded the neuronal calcium activity in somatosensory cortex regardless of neuronal subtypes. Detailed dissection of FUS-mediated modulation of different neuronal subtypes, with systematic analysis of mechanosensitive channel expression profiles in these neurons will be helpful but is beyond the scope of this study. Moreover, it was reported that FUS could evoke region-specific neuronal response profiles in awake mammalian brain^65^. Therefore, the suppression effect of FUS at 0.521MHz fundamental frequency and 0.91 W/cm^2^ intensity in somatosensory cortical neurons observed in our study might not be applicable to other brain regions.

The different effects of FUS on cortical astrocyte and neuronal calcium dynamics suggest that these two cell types might respond to FUS through distinct mechanisms. Given the critical roles of astrocyte in synaptic transmission and neurotransmitter regulation^16,66,67^, it is possible that FUS first activates astrocytes then modulates neuronal activity indirectly. Neuronal-astrocyte co-culture experiments also supported that FUS could induce TRPA1-mediated calcium increase in astrocytes to promote glutamate or GABA signaling in neurons^17,18^. Whether the suppression of neuronal calcium activity observed in our study was caused by FUS-mediated activation of astrocyte-dependent inhibitory neurotransmitter release requires further investigation.

We carefully controlled heat effects during FUS stimulation and 2P imaging. While mild transient temperature increases were associated with two-photon laser excitation, no significant temperature changes were detected during FUS stimulation, confirming that the observed effects were independent of thermal effects of FUS. For 2P imaging, however, it was reported that infrared excitation could induce heating and calcium microdomain hyperactivity in cortical astrocytes^68,69^. Although we minimized the laser power while still maintaining necessary spatiotemporal resolution, we still observed an increase of calcium activity in astrocyte subregions at the end of 2P image recording, but not neurons, especially in anesthetized mice in the absence of FUS treatment. Nevertheless, the percentage of astrocyte calcium activity induced by FUS at 1.5 W/cm^2^ intensity was significantly higher than the baseline increase that was induced by 2P imaging alone, indicating a profound effect of FUS on astrocyte calcium dynamics. These results also suggest that careful monitoring of the temperature in the brain during FUS treatment and 2P imaging is important for the validation of FUS-dependent neuromodulation.

An important consideration in transcranial ultrasound research is the possibility of stimulation of the auditory system^70–72^. Instead of a rectangular waveform, our FUS stimulations were set in pulsed and smoothed waveforms to eliminate the auditory confound that could mask the direct FUS modulation^73^. While we did not observe overt behavioral reactions in our mice, we cannot exclude the possibility that some component of the neuronal or astrocytic changes observed here was partially mediated by skull vibrations or harmonic components. Future experiments with pharmacological blockade of cochlear activity or inducible deafening of mice^74^ could help verify the direct effects of FUS on astrocyte and neuronal calcium dynamics. Another limitation of our study is that we did not record neuronal and astrocyte calcium transients simultaneously. Multiple technical and biological constraints informed this choice. First, simultaneous 2P imaging of two sets of genetically encoded calcium indicators (e.g., GCaMP for astrocytes and jRGECO for neurons) requires spectral unmixing or separate emission filters, which can lower signal-to-noise ratios and compromise temporal resolution if done simultaneously. Importantly, the extended exposure to dual infrared laser would induce significant temperature rise in mouse cortex. To keep the temperature rise below 1 ℃ during 2P imaging and minimize its effect on baseline calcium activity, we chose to image astrocytes and neurons separately. Second, astrocyte calcium transients typically exhibit slower kinetics and broader spatial domains, whereas neuronal signals are more rapid and localized^75–77^. Optimizing acquisition settings (frame rate, field of view) for both cell types can require significant tradeoffs in data quality, especially for single-cell or microdomain tracing. Despite these limitations, future work using optimized multiplex imaging strategies will be necessary to understand precisely how astrocytes and neurons interact during FUS neuromodulation in real time.

In summary, we present the experimental results acquired by simultaneous 2P calcium imaging and FUS stimulation. The calcium imaging data suggests that FUS suppressed neuronal calcium activity but promoted calcium responses in astrocyte microdomains in somatosensory cortex of the awake mice. Additionally, FUS also increased the calcium transients in astrocytes in anesthetized mice. These results indicate the potential distinct mechanisms for FUS-dependent modulation of neurons and astrocytes. The ability of FUS to selectively suppress neuronal activity while enhancing astrocytic responses in awake animals opens exciting possibilities for its application in treating neurological disorders characterized by hyperactive neuronal circuits or impaired astrocytic function, such as Alzheimer’s disease, epilepsy and neuropathic pain^29,78,79^. Future studies are warranted to investigate the underlying mechanisms driving these effects, such as potential contributions of mechanosensitive ion channels or downstream signaling pathways. Furthermore, the development of targeted FUS paradigms optimized for specific cell types or brain states could further enhance its therapeutic application.

## CRediT authorship contribution statement

**Yu Yong:** Conceptualization, Methodology, Investigation, Data acquisition and analysis, Writing, Review and editing. **Hao Jiang:** Methodology, Investigation, Data analysis, Writing, Review and editing. **Chaofeng Qiao:** Data acquisition. **Zhuoyan Liu:** Methodology and Data acquisition. **Xiaohan Zhou:** Data analysis. **Yufeng Zhou:** Methodology. **Fenfang Li:** Conceptualization, Methodology, Investigation, Supervision, Writing, Review and editing, Funding acquisition.

## Acknowledgement

This work was funded by the National Natural Science Foundation of China (No. 12204322) and the Natural Science Foundation of Guangdong Province (Nos. 2023A1515010649). We thank the technical support of the core facility of Shenzhen Bay Laboratory for mice maintenance and two-photon microscopy. We thank Prof. Wenbiao Gan (Shenzhen Bay Laboratory Neurological Institute) and Dr. Zhihai Qiu (Guangdong Institute of Intelligence Science and Technology) for valuable comments and discussion. The authors acknowledge the assistance of Jianpeng Wei and Xueshuo Guo for the characterization of the acoustic pressure field of the ultrasound transducer and laser power measurement, respectively.

## Data availability

The datasets generated during and/or analyzed during the current study are available from the corresponding author on reasonable request. The code is available at GitHub: https://github.com/Johnsun4211/Neu-ASTGCa-data-analysis

## References

1 Lewis, P. M., Thomson, R. H., Rosenfeld, J. V. & Fitzgerald, P. B. Brain Neuromodulation Techniques: A Review. Neuroscientist 22, 406–421 (2016). 10.1177/1073858416646707

2 Davidson, B. et al. Neuromodulation techniques - From non-invasive brain stimulation to deep brain stimulation. Neurotherapeutics 21, e00330 (2024). 10.1016/j.neurot.2024.e00330

3 Nguyen, J. P., Nizard, J., Keravel, Y. & Lefaucheur, J. P. Invasive brain stimulation for the treatment of neuropathic pain. Nat Rev Neurol 7, 699–709 (2011). 10.1038/nrneurol.2011.138

4 Reithler, J., Peters, J. C. & Sack, A. T. Multimodal transcranial magnetic stimulation: using concurrent neuroimaging to reveal the neural network dynamics of noninvasive brain stimulation. Prog Neurobiol 94, 149–165 (2011). 10.1016/j.pneurobio.2011.04.004

5 Weise, K. et al. Precise motor mapping with transcranial magnetic stimulation. Nat Protoc 18, 293–318 (2023). 10.1038/s41596-022-00776-6

6 Woods, A. J. et al. A technical guide to tDCS, and related non-invasive brain stimulation tools. Clin Neurophysiol 127, 1031–1048 (2016). 10.1016/j.clinph.2015.11.012

7 Rabut, C. et al. Ultrasound Technologies for Imaging and Modulating Neural Activity. Neuron 108, 93–110 (2020). 10.1016/j.neuron.2020.09.003

8 Kuhn, T. et al. Transcranial focused ultrasound selectively increases perfusion and modulates functional connectivity of deep brain regions in humans. Front Neural Circuits 17, 1120410 (2023). 10.3389/fncir.2023.1120410

9 Tufail, Y. et al. Transcranial pulsed ultrasound stimulates intact brain circuits. Neuron 66, 681–694 (2010). 10.1016/j.neuron.2010.05.008

10 Folloni, D. et al. Manipulation of Subcortical and Deep Cortical Activity in the Primate Brain Using Transcranial Focused Ultrasound Stimulation. Neuron 101, 1109–1116 e1105 (2019). 10.1016/j.neuron.2019.01.019

11 Yaakub, S. N. et al. Transcranial focused ultrasound-mediated neurochemical and functional connectivity changes in deep cortical regions in humans. Nat Commun 14, 5318 (2023). 10.1038/s41467-023-40998-0

12 Fomenko, A., Neudorfer, C., Dallapiazza, R. F., Kalia, S. K. & Lozano, A. M. Low-intensity ultrasound neuromodulation: An overview of mechanisms and emerging human applications. Brain Stimul 11, 1209–1217 (2018). 10.1016/j.brs.2018.08.013

13 Prieto, M. L., Firouzi, K., Khuri-Yakub, B. T., Madison, D. V. & Maduke, M. Spike frequency-dependent inhibition and excitation of neural activity by high-frequency ultrasound. J Gen Physiol 152 (2020). 10.1085/jgp.202012672

14 Naour, A. L. et al. Do astrocytes respond to light, sound, or electrical stimulation? Neural Regen Res 18, 2343–2347 (2023). 10.4103/1673-5374.371343

15 Bazargani, N. & Attwell, D. Astrocyte calcium signaling: the third wave. Nat Neurosci 19, 182–189 (2016). 10.1038/nn.4201

16 Oliveira, J. F. & Araque, A. Astrocyte regulation of neural circuit activity and network states. Glia 70, 1455–1466 (2022). 10.1002/glia.24178

17 Oh, S. J. et al. Ultrasonic Neuromodulation via Astrocytic TRPA1. Curr Biol 29, 3386–3401 e3388 (2019). 10.1016/j.cub.2019.08.021

18 Mishima, T. et al. Repetitive pulsed-wave ultrasound stimulation suppresses neural activity by modulating ambient GABA levels via effects on astrocytes. Front Cell Neurosci 18, 1361242 (2024). 10.3389/fncel.2024.1361242

19 Lee, K. et al. Ultrasonocoverslip: In-vitro platform for high-throughput assay of cell type-specific neuromodulation with ultra-low-intensity ultrasound stimulation. Brain Stimul 16, 1533–1548 (2023). 10.1016/j.brs.2023.08.002

20 Deng, Z. et al. Ultrasound-mediated augmented exosome release from astrocytes alleviates amyloid-beta-induced neurotoxicity. Theranostics 11, 4351–4362 (2021). 10.7150/thno.52436

21 Liu, S. H., Lai, Y. L., Chen, B. L. & Yang, F. Y. Ultrasound Enhances the Expression of Brain-Derived Neurotrophic Factor in Astrocyte Through Activation of TrkB-Akt and Calcium-CaMK Signaling Pathways. Cereb Cortex 27, 3152–3160 (2017). 10.1093/cercor/bhw169

22 Yang, F. Y., Lu, W. W., Lin, W. T., Chang, C. W. & Huang, S. L. Enhancement of Neurotrophic Factors in Astrocyte for Neuroprotective Effects in Brain Disorders Using Low-intensity Pulsed Ultrasound Stimulation. Brain Stimul 8, 465–473 (2015). 10.1016/j.brs.2014.11.017

23 Santello, M., Toni, N. & Volterra, A. Astrocyte function from information processing to cognition and cognitive impairment. Nat Neurosci 22, 154–166 (2019). 10.1038/s41593-018-0325-8

24 Zhang, Y. V., Ormerod, K. G. & Littleton, J. T. Astrocyte Ca(2+) Influx Negatively Regulates Neuronal Activity. eNeuro 4 (2017). 10.1523/ENEURO.0340-16.2017

25 Pascual, O. et al. Astrocytic purinergic signaling coordinates synaptic networks. Science 310, 113–116 (2005). 10.1126/science.1116916

26 Chen, J. et al. Heterosynaptic long-term depression mediated by ATP released from astrocytes. Glia 61, 178–191 (2013). 10.1002/glia.22425

27 Durkee, C., Kofuji, P., Navarrete, M. & Araque, A. Astrocyte and neuron cooperation in long-term depression. Trends Neurosci 44, 837–848 (2021). 10.1016/j.tins.2021.07.004

28 Cheng, Y. T. et al. Social deprivation induces astrocytic TRPA1-GABA suppression of hippocampal circuits. Neuron 111, 1301–1315 e1305 (2023). 10.1016/j.neuron.2023.01.015

29 Shah, D. et al. Astrocyte calcium dysfunction causes early network hyperactivity in Alzheimer’s disease. Cell Rep 40, 111280 (2022). 10.1016/j.celrep.2022.111280

30 Xu, R. S., Wu, X. M. & Xiong, Z. Q. Low-intensity ultrasound directly modulates neural activity of the cerebellar cortex. Brain Stimul 16, 918–926 (2023). 10.1016/j.brs.2023.05.012

31 Wang, X., Zhang, Y., Zhang, K. & Yuan, Y. Influence of behavioral state on the neuromodulatory effect of low-intensity transcranial ultrasound stimulation on hippocampal CA1 in mouse. Neuroimage 241, 118441 (2021). 10.1016/j.neuroimage.2021.118441

32 Tripathi, K. et al. Direct Activation of Cortical Neurons in the Primary Somatosensory Cortex of the Rat in Vivo Using Focused Ultrasound. Ultrasound Med Biol 46, 2349–2360 (2020). 10.1016/j.ultrasmedbio.2020.06.003

33 Murphy, K. R. et al. A tool for monitoring cell type-specific focused ultrasound neuromodulation and control of chronic epilepsy. Proc Natl Acad Sci U S A 119, e2206828119 (2022). 10.1073/pnas.2206828119

34 Mesik, L. et al. Transcranial Low-Intensity Focused Ultrasound Stimulation of the Visual Thalamus Produces Long-Term Depression of Thalamocortical Synapses in the Adult Visual Cortex. J Neurosci 44 (2024). 10.1523/JNEUROSCI.0784-23.2024

35 Darrow, D. P., O’Brien, P., Richner, T. J., Netoff, T. I. & Ebbini, E. S. Reversible neuroinhibition by focused ultrasound is mediated by a thermal mechanism. Brain Stimul 12, 1439–1447 (2019). 10.1016/j.brs.2019.07.015

36 Cheng, Z. et al. High resolution ultrasonic neural modulation observed via in vivo two-photon calcium imaging. Brain Stimul 15, 190–196 (2022). 10.1016/j.brs.2021.12.005

37 Choi, T. et al. Bidirectional Neuronal Control of Epileptiform Activity by Repetitive Transcranial Focused Ultrasound Stimulations. Adv Sci (Weinh) 11, e2302404 (2024). 10.1002/advs.202302404

38 Zhang, T. et al. Excitatory-inhibitory modulation of transcranial focus ultrasound stimulation on human motor cortex. CNS Neurosci Ther 29, 3829–3841 (2023). 10.1111/cns.14303

39 Yang, P. F. et al. Differential dose responses of transcranial focused ultrasound at brain regions indicate causal interactions. Brain Stimul 15, 1552–1564 (2022). 10.1016/j.brs.2022.12.003

40 Yang, P. F. et al. Bidirectional and state-dependent modulation of brain activity by transcranial focused ultrasound in non-human primates. Brain Stimul 14, 261–272 (2021). 10.1016/j.brs.2021.01.006

41 Murphy, K. R. et al. Optimized ultrasound neuromodulation for non-invasive control of behavior and physiology. Neuron 112, 3252–3266 e3255 (2024). 10.1016/j.neuron.2024.07.002

42 Giovannucci, A. et al. CaImAn an open source tool for scalable calcium imaging data analysis. Elife 8 (2019). 10.7554/eLife.38173

43 Wang, Y. et al. Accurate quantification of astrocyte and neurotransmitter fluorescence dynamics for single-cell and population-level physiology. Nat Neurosci 22, 1936–1944 (2019). 10.1038/s41593-019-0492-2

44 Mi, X. et al. Fast, Accurate, and Versatile Data Analysis Platform for the Quantification of Molecular Spatiotemporal Signals. bioRxiv (2024). 10.1101/2024.05.02.592259

45 Tsunematsu, T., Sakata, S., Sanagi, T., Tanaka, K. F. & Matsui, K. Region-Specific and State-Dependent Astrocyte Ca(2+) Dynamics during the Sleep-Wake Cycle in Mice. J Neurosci 41, 5440–5452 (2021). 10.1523/JNEUROSCI.2912-20.2021

46 Poskanzer, K. E. & Yuste, R. Astrocytes regulate cortical state switching in vivo. Proc Natl Acad Sci U S A 113, E2675–2684 (2016). 10.1073/pnas.1520759113

47 Srinivasan, R. et al. Ca(2+) signaling in astrocytes from Ip3r2(-/-) mice in brain slices and during startle responses in vivo. Nat Neurosci 18, 708–717 (2015). 10.1038/nn.4001

48 Yuan, Y. et al. The Effect of Low-Intensity Transcranial Ultrasound Stimulation on Neural Oscillation and Hemodynamics in the Mouse Visual Cortex Depends on Anesthesia Level and Ultrasound Intensity. IEEE Trans Biomed Eng 68, 1619–1626 (2021). 10.1109/TBME.2021.3050797

49 Thrane, A. S. et al. General anesthesia selectively disrupts astrocyte calcium signaling in the awake mouse cortex. Proc Natl Acad Sci U S A 109, 18974–18979 (2012). 10.1073/pnas.1209448109

50 Bojarskaite, L. et al. Astrocytic Ca(2+) signaling is reduced during sleep and is involved in the regulation of slow wave sleep. Nat Commun 11, 3240 (2020). 10.1038/s41467-020-17062-2

51 Jiwaji, Z. et al. General anesthesia alters CNS and astrocyte expression of activity-dependent and activity-independent genes. Front Netw Physiol 3, 1216366 (2023). 10.3389/fnetp.2023.1216366

52 Ye, P. P., Brown, J. R. & Pauly, K. B. Frequency Dependence of Ultrasound Neurostimulation in the Mouse Brain. Ultrasound Med Biol 42, 1512–1530 (2016). 10.1016/j.ultrasmedbio.2016.02.012

53 Kubanek, J., Shukla, P., Das, A., Baccus, S. A. & Goodman, M. B. Ultrasound Elicits Behavioral Responses through Mechanical Effects on Neurons and Ion Channels in a Simple Nervous System. J Neurosci 38, 3081–3091 (2018). 10.1523/JNEUROSCI.1458-17.2018

54 Yu, K., Niu, X., Krook-Magnuson, E. & He, B. Intrinsic functional neuron-type selectivity of transcranial focused ultrasound neuromodulation. Nat Commun 12, 2519 (2021). 10.1038/s41467-021-22743-7

55 Newman, M. et al. Ultrasound Modulates Calcium Activity in Cultured Neurons, Glial Cells, Endothelial Cells and Pericytes. Ultrasound Med Biol 50, 341–351 (2024). 10.1016/j.ultrasmedbio.2023.11.004

56 Zhu, J. et al. The mechanosensitive ion channel Piezo1 contributes to ultrasound neuromodulation. Proc Natl Acad Sci U S A 120, e2300291120 (2023). 10.1073/pnas.2300291120

57 Oh, S. J. et al. Ultrasonic Neuromodulation via Astrocytic TRPA1. Curr Biol 30, 948 (2020). 10.1016/j.cub.2020.02.042

58 Liao, D., Li, F., Lu, D. & Zhong, P. Activation of Piezo1 mechanosensitive ion channel in HEK293T cells by 30 MHz vertically deployed surface acoustic waves. Biochem Biophys Res Commun 518, 541–547 (2019). 10.1016/j.bbrc.2019.08.078

59 Yoo, S., Mittelstein, D. R., Hurt, R. C., Lacroix, J. & Shapiro, M. G. Focused ultrasound excites cortical neurons via mechanosensitive calcium accumulation and ion channel amplification. Nat Commun 13, 493 (2022). 10.1038/s41467-022-28040-1

60 Li, D. et al. Low-Intensity Pulsed Ultrasound Dynamically Modulates the Migration of BV2 Microglia. Ultrasound Med Biol 51, 494–507 (2025). 10.1016/j.ultrasmedbio.2024.11.010

61 Li, F. et al. Mechanically induced integrin ligation mediates intracellular calcium signaling with single pulsating cavitation bubbles. Theranostics 11, 6090–6104 (2021). 10.7150/thno.56813

62 Sitdikova, G. et al. Isoflurane suppresses early cortical activity. Ann Clin Transl Neurol 1, 15–26 (2014). 10.1002/acn3.16

63 Hentschke, H., Raz, A., Krause, B. M., Murphy, C. A. & Banks, M. I. Disruption of cortical network activity by the general anaesthetic isoflurane. Br J Anaesth 119, 685–696 (2017). 10.1093/bja/aex199

64 Sherman, J. et al. Ultrasound pulse repetition frequency preferentially activates different neuron populations independent of cell type. J Neural Eng 21 (2024). 10.1088/1741-2552/ad731c

65 Tseng, H. A. et al. Region-specific effects of ultrasound on individual neurons in the awake mammalian brain. iScience 24, 102955 (2021). 10.1016/j.isci.2021.102955

66 Halassa, M. M. & Haydon, P. G. Integrated brain circuits: astrocytic networks modulate neuronal activity and behavior. Annu Rev Physiol 72, 335–355 (2010). 10.1146/annurev-physiol-021909-135843

67 Parpura, V. & Haydon, P. G. Physiological astrocytic calcium levels stimulate glutamate release to modulate adjacent neurons. Proc Natl Acad Sci U S A 97, 8629–8634 (2000). 10.1073/pnas.97.15.8629

68 Podgorski, K. & Ranganathan, G. Brain heating induced by near-infrared lasers during multiphoton microscopy. J Neurophysiol 116, 1012–1023 (2016). 10.1152/jn.00275.2016

69 Schmidt, E. & Oheim, M. Infrared Excitation Induces Heating and Calcium Microdomain Hyperactivity in Cortical Astrocytes. Biophys J 119, 2153–2165 (2020). 10.1016/j.bpj.2020.10.027

70 Qi, X. et al. Low-Intensity Ultrasound Causes Direct Excitation of Auditory Cortical Neurons. Neural Plast 2021, 8855055 (2021). 10.1155/2021/8855055

71 Guo, H. et al. Ultrasound Produces Extensive Brain Activation via a Cochlear Pathway. Neuron 98, 1020–1030 e1024 (2018). 10.1016/j.neuron.2018.04.036

72 Sato, T., Shapiro, M. G. & Tsao, D. Y. Ultrasonic Neuromodulation Causes Widespread Cortical Activation via an Indirect Auditory Mechanism. Neuron 98, 1031–1041 e1035 (2018). 10.1016/j.neuron.2018.05.009

73 Mohammadjavadi, M. et al. Elimination of peripheral auditory pathway activation does not affect motor responses from ultrasound neuromodulation. Brain Stimul 12, 901–910 (2019). 10.1016/j.brs.2019.03.005

74 Guo, H. et al. Effects of focused ultrasound in a “clean” mouse model of ultrasonic neuromodulation. iScience 26, 108372 (2023). 10.1016/j.isci.2023.108372

75 Shigetomi, E., Patel, S. & Khakh, B. S. Probing the Complexities of Astrocyte Calcium Signaling. Trends Cell Biol 26, 300–312 (2016). 10.1016/j.tcb.2016.01.003

76 Lines, J. et al. A spatial threshold for astrocyte calcium surge. Elife 12 (2024). 10.7554/eLife.90046

77 Winship, I. R., Plaa, N. & Murphy, T. H. Rapid astrocyte calcium signals correlate with neuronal activity and onset of the hemodynamic response in vivo. J Neurosci 27, 6268–6272 (2007). 10.1523/JNEUROSCI.4801-06.2007

78 Purnell, B. S., Alves, M. & Boison, D. Astrocyte-neuron circuits in epilepsy. Neurobiol Dis 179, 106058 (2023). 10.1016/j.nbd.2023.106058

79 Takeda, I. et al. Controlled activation of cortical astrocytes modulates neuropathic pain-like behaviour. Nat Commun 13, 4100 (2022). 10.1038/s41467-022-31773-8

