## Supplementary Information for "Ultrasonic Modulation of Astrocytic and Neuronal Calcium Dynamics in Mouse Cortex"

### Supplemental Materials

#### 1. Acoustic field measurement

To measure the output power of ultrasound transducer for *in vivo* studies, a submersible hydrophone (HNR0500, ONDA) was placed beneath the coverslip used for cranial window in a water tank. The hydrophone was positioned using a 3D manipulator controlled by a $C^{++}$ interface (SoundFieldScanning.cpp), aligned to the geometric center of the transducer. For spatial mapping, the scanning protocol involved discretizing designated XY, XZ planes into grid matrices within predefined spatial ranges. At each grid node, the hydrophone recorded acoustic pressure, frequency, and waveform parameters. To identify the acoustic focal region, XZ-plane and XY-plane scans were initially performed. A high-resolution XY-plane scan (step size: 0.2 mm) was subsequently conducted within the focal plane to capture detailed field distributions, followed by acoustic pressure measurements at the focal point under varying input voltages and pressure calibration. The acquired time-domain signals, including peak-to-peak amplitudes, were post-processed using MATLAB (SoundFieldAnalysis.m) for noise filtering, signal averaging, and 3D heatmap visualization. This methodology enabled comprehensive spatial reconstruction of the acoustic field, facilitating quantitative analysis of pressure gradients and focal zone characteristics.

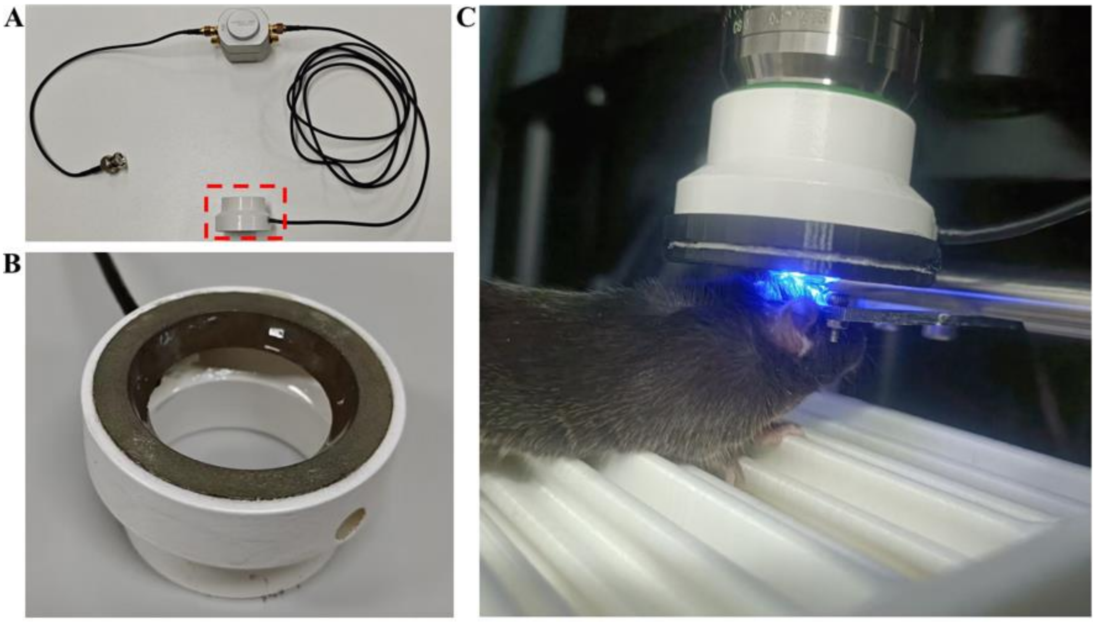


Fig. S1. Ultrasound transducer and experimental setup. (A) A photo of the custom-built 0.521 MHz ring-shaped ultrasound transducer. (B) Enlarged view of the dashed box in panel A showing the emitting surface of the transducer with a concentric spherical curvature. (C) A photo showing the transducer with a coupling cone filled with de-gassed ultrasound gel installed on the Olympus 25× objective for *in vivo* two-photon imaging of mice cortex.

**2.Calculation of total events area**

This computational procedure was implemented through the custom MATLAB script UniqueEvents.m. The workflow consisted of: (1) selecting the target AQuA file and specifying its directory path; (2) setting a breakpoint at the predefined debugging position; (3) executing the script to initialize the graphical user interface (GUI). Within the GUI environment, a grayscale threshold was empirically determined to segment cellular regions in time-averaged video frames, while activity-responsive regions were automatically identified through AQuA data parsing. The total event area was subsequently derived by computing the union of these two regions sets.

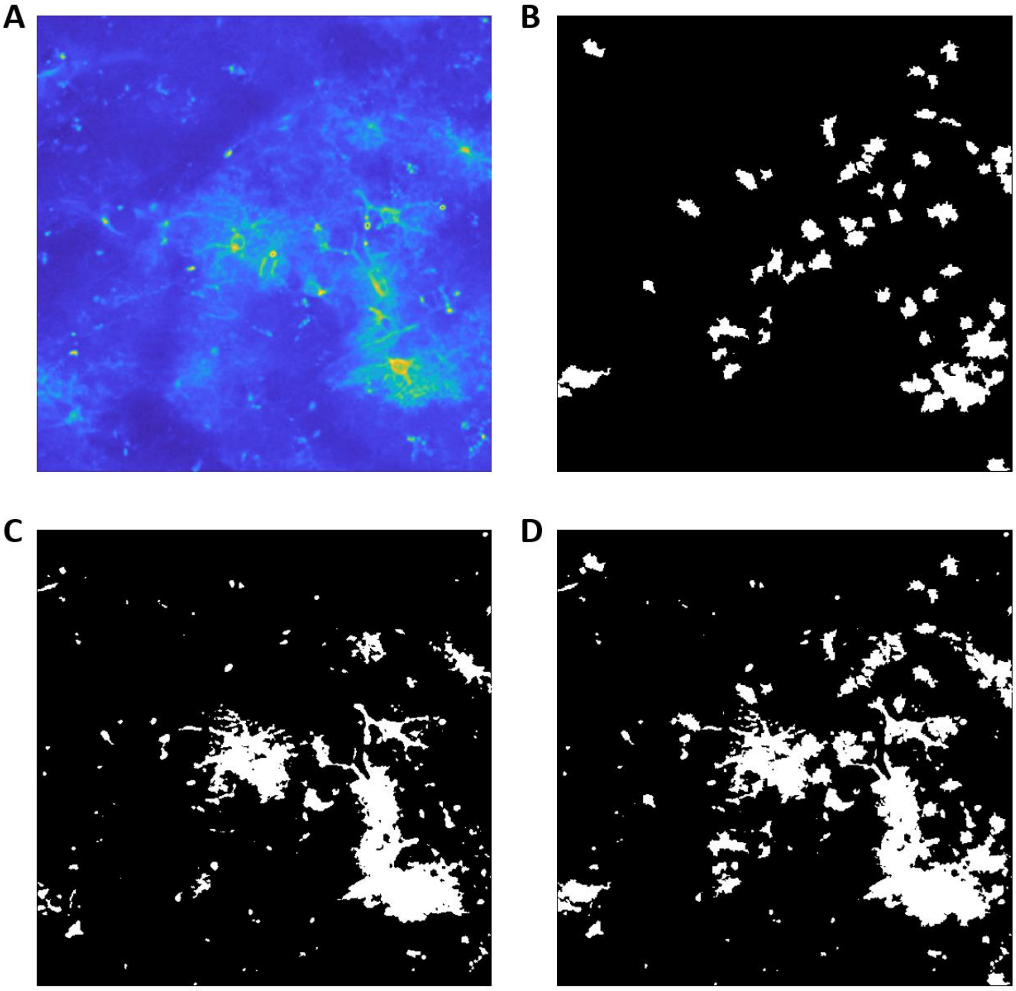


Fig. S2. Computational workflow for total event area determination. (A) Pseudo-colored representation of time-averaged intensity projections. (B) Binary mask identifying regions of activity (ROA) through AquA2. (C) Cellular region segmentation via grayscale thresholding applied to Panel A. (D) Morphological union operation integrating regions from B and C.

**3. Calcium activity functional clustering analysis**

**AUC** K-means Clustering Analysis: The AUC data for astrocytes or neurons under different ultrasound stimulation conditions were imported into a custom Python program for clustering analysis (dffAnalysis_Cluster.py). The 'kmeans.inertia_' function was used to evaluate clustering performance, and an elbow plot was generated to determine the optimal number of clusters. K-means clustering was then performed on the entire dataset to qualitatively assess calcium signal patterns. The clustered (annotated) data were used for further statistical analysis.

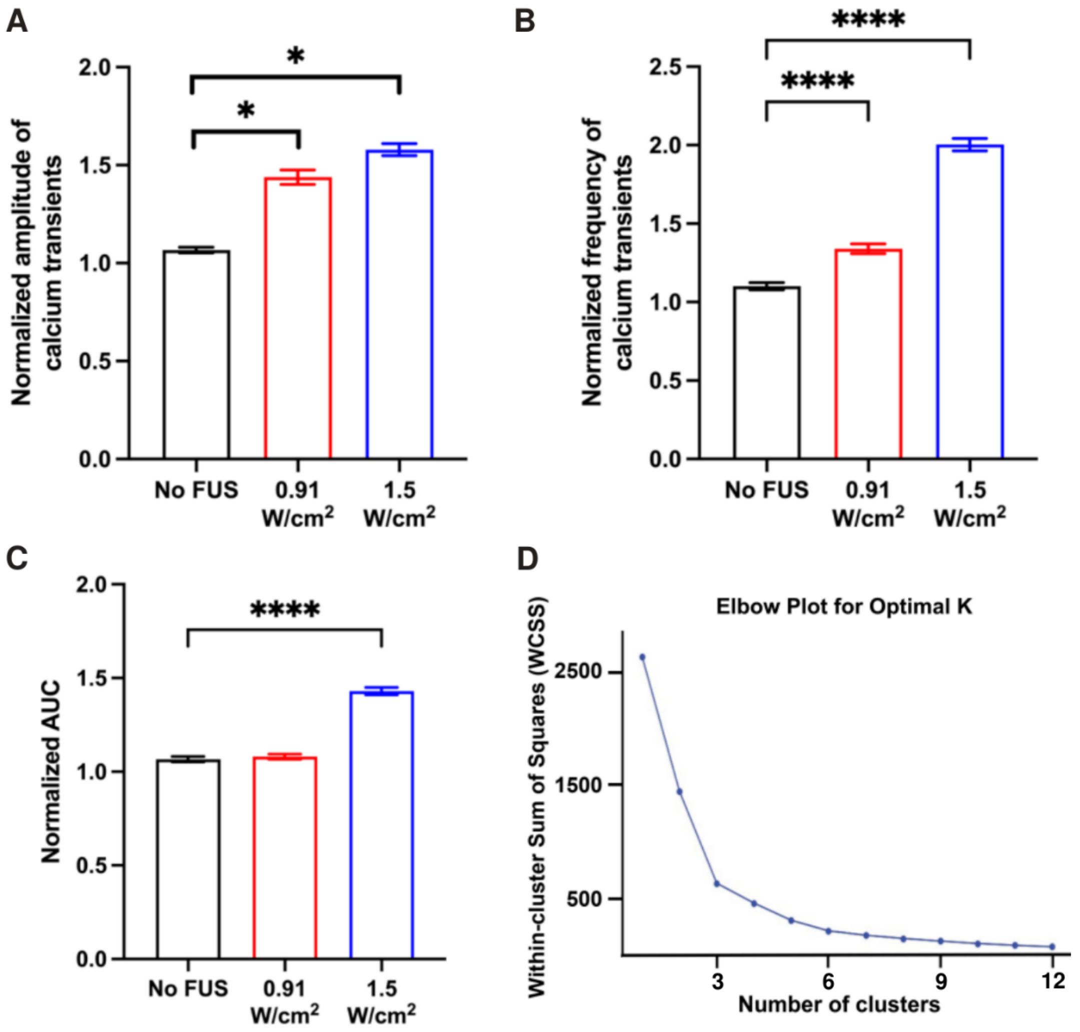


**Figure S3: Analysis of astrocyte calcium activity during FUS and functional clustering in awake mice.** (A) Normalized amplitude of calcium transients, (B) normalized frequency of calcium transients, and (C) normalized area under the curve (AUC) of astrocyte calcium responses during FUS stimulation at intensities of 0 (No FUS), 0.91, or 1.5 W/cm$^2$ in awake mice, respectively. The same dataset was presented in Figure 2. *$p$<0.05, ****$p$<0.0001. (D) The elbow plot for optimal number of clusters used in astrocyte calcium K-means analysis.

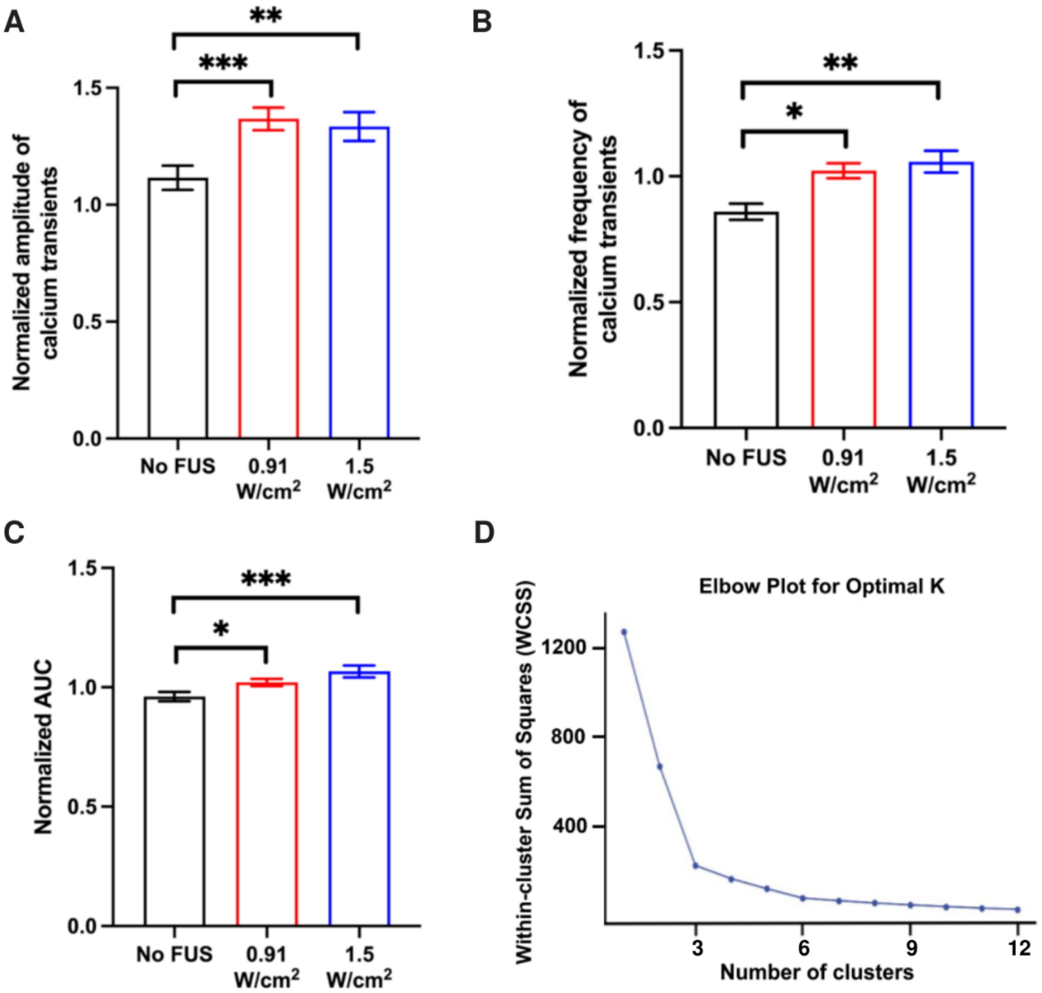


**Figure S4: Analysis of astrocyte calcium activity during FUS and functional clustering in lightly anesthetized mice.** (A) Normalized amplitude of calcium transients, (B) normalized frequency of calcium transients, and (C) normalized area under the curve (AUC) of astrocyte calcium responses during FUS stimulation at intensities of 0 (No FUS), 0.91, or 1.5 W/cm$^2$ in lightly anesthetized mice, respectively. The same dataset was presented in Figure 4. *$p<0.05$, ** $p<0.01$, *** $p<0.001$, ****$p<0.0001$. (D) The elbow plot for optimal number of clusters used in astrocyte calcium K-means analysis.

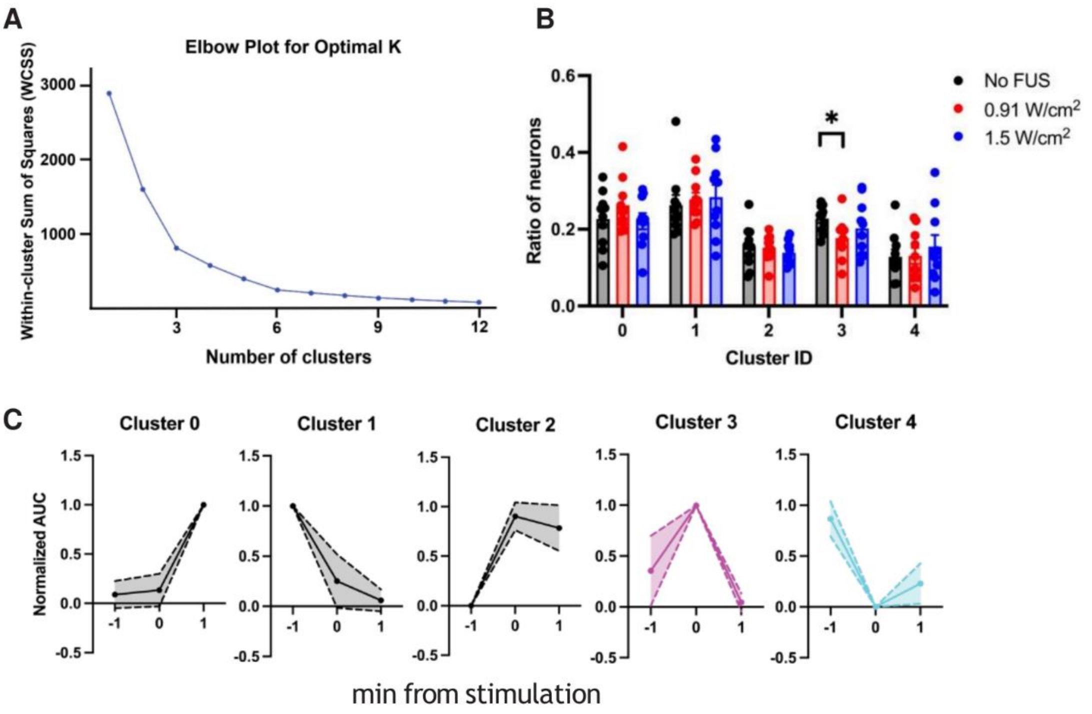


**Figure S5: Neuronal calcium activity functional clustering analysis in awake mice.** (A) The elbow plot for optimal number of clusters used in K-means analysis. (B) The average response profiles (solid lines) with SD labeled by shadow area for each cluster are shown in the bottom panel. The cluster 3 (magenta) and cluster 4 (cyan) are identified as activated and inhibited neuronal calcium response during FUS, respectively. The dataset was also presented in Figure 7B. (C) Ratio of neurons for each calcium activity functional subpopulation in awake mice untreated and treated with ultrasound stimulation. Data are presented as mean ± SEM. $n$ = 10 mice for each condition. Significant differences were determined by two-way ANOVA with Dunnett's multiple comparisons test. *$p$<0.05.

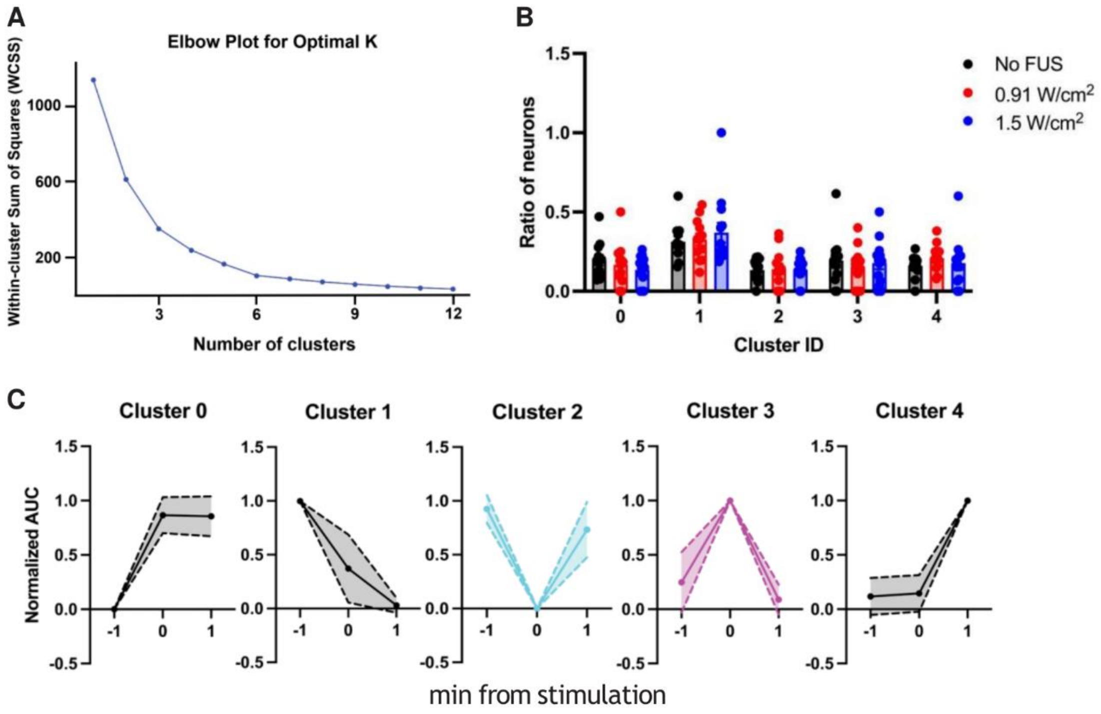


**Figure S6: Neuronal calcium activity functional clustering analysis in lightly anesthetized mice.** (A) The elbow plot for optimal number of clusters used in K-means analysis. (B) The average response profiles (solid lines) with SD labeled by shadow area for each cluster are shown in the bottom panel. Cluster 2 (cyan) and cluster 3 (magenta) are identified as inhibited and activated neuronal calcium response during FUS, respectively. The dataset was also presented in Figure 9B. (C) Ratio of neurons for each calcium activity functional subpopulation in lightly anesthetized mice untreated and treated with ultrasound stimulation. Data are presented as mean ± SEM. $n$ = 13 mice for each condition.
